# CHARIOT-AAV: Conjugation of diverse vectors to adeno-associated viruses for delivery of large genes

**DOI:** 10.64898/2026.04.13.718054

**Authors:** Keisuke Nagao, Tatsuya Osaki, Sho Toyonaga, Chae Gyu Lee, Emmanuel Vargas Paniagua, Emily Crespin Guerra, Scott Machen, Jacob L. Beckham, Mriganka Sur, Daniel Griffith Anderson, Robert J. Macfarlane, Polina Anikeeva

## Abstract

Systemic, tissue-specific delivery of large transgenes exceeding the packaging capacity of adeno-associated viruses (AAVs) remains a key translational challenge for molecular therapeutics. Vectors with larger capacities, such as lentiviral vectors (LVVs) and lipid nanoparticles (LNPs), often lack adjustable, tissue-specific tropisms. Here we report CHARIOT-AAV (Crosslinked Hybrid Architectures for Robust, Interchangeable, and Organ-specific Targeting with AAV), a platform where diverse delivery vectors are conjugated to AAVs, thereby achieving tissue-specific tropism of AAVs and expanded cargo capacity. AAV-AAV conjugates packaging split SpCas9 constructs in AAV.CAP-B10 capsids demonstrate a ∼2-fold increase in brain gene editing efficiency over unconjugated AAV cocktails after intravenous injection. In addition to AAV-AAV conjugates, AAV-LVV and AAV-LNP conjugates achieve AAV-guided delivery of genetic payloads to target cells. Furthermore, AAV-LNP conjugates enable systemic delivery of mRNAs to brain endothelial cells. CHARIOT-AAV thus provides a modular platform for systemic, tissue-specific delivery of diverse therapeutics beyond the limits of individual vectors.

## Main

Systemic delivery of genetic cargo to target tissues via intravenous injection could transform molecular therapies for genetic disorders and empower research with genetic models, yet it remains a fundamental challenge^1,2^. Among delivery platforms, recombinant adeno-associated viruses (AAVs) are one of the most promising vehicles, offering low pathogenicity, limited immunogenicity, and tunable tissue-specific tropism^3–11^. However, AAV’s gene packaging capacity is limited to ∼4.8 kb^12^, which must accommodate not only molecular tools but also regulatory elements such as cell-type-specific promoters and organelle trafficking signals. Transformative molecular tools such as CRISPR-Cas gene editing systems exceed this capacity (> ∼6 kb)^1^, and even smaller functional equivalents (e.g. CasMINI^13^, NanoCas^14^) that fit within a single capsid leave insufficient room for the long regulatory sequences necessary for cell-type specificity. Alternative vehicles such as lentiviral vectors (LVVs) and lipid nanoparticles (LNPs) can accommodate larger cargo (∼10 kb)^15–17^; however, these vectors offer limited tissue specificity compared to AAVs^18,19^. There remains a need for platforms that accommodate large and diverse genetic cargo while achieving tissue specificity comparable to that of AAVs.

To circumvent AAV’s packaging limit, previous work has pursued dual-AAV strategies that split a large transgene across two vectors, relying on subsequent reconstitution of the full-length gene or protein in target cells^20,21^. However, these approaches demand simultaneous transduction of both constructs packaged in separate AAVs, a requirement that potentially compromises overall efficacy^22,23^ and necessitates elevated vector doses that raise safety concerns^22,24^. Meanwhile, non-AAV vectors, while promising^25–27^, are yet to achieve tissue-specific tropism and versatility comparable to AAVs.

To enable systemic, tissue-specific delivery of payloads beyond the packaging capacity of AAVs, we developed a platform that employs surface chemistry to covalently couple AAV capsids to diverse molecular delivery vehicles, resulting in oligomeric constructs that recapitulate the tissue-specific tropism of AAVs while expanding cargo capacity. We term this platform CHARIOT-AAV (Crosslinked Hybrid Architectures for Robust, Interchangeable, and Organ-specific Targeting with AAV). Using this platform, we first address the co-delivery bottleneck inherent in dual-AAV strategies by conjugating multiple AAV capsids to ensure simultaneous transduction of split transgenes (**Fig. 1a**). The presence of *Lys* residues on the surfaces of AAV capsids permits the chemical modification with complementary tetrazine (Tz) or *trans*-cyclooctene (TCO) moieties via facile carbodiimide chemistry, enabling their crosslinking via inverse electron-demand Diels-Alder (IEDDA) click chemistry^28^. We applied this approach to create conjugates of two AAV serotypes: AAV-DJ (AAV_DJ_), which efficiently transduces HEK293T cells, and AAV9, which does not (**Fig. 1b**, Fig. S1)^29^. Each capsid packaged a distinct fluorescent reporter, NLS-GFP (nuclear localization signal-tagged green fluorescent protein) in AAV9 and a red-emitting fluorescent protein mRuby2 in AAV_DJ_, under a ubiquitous CAG promoter. In cultures transduced with a 1:1 cocktail of AAV9-CAG::NLS-GFP and AAV_DJ_-CAG::mRuby2, negligible expression of NLS-GFP was observed, while mRuby2 exhibited strong fluorescence (**Fig. 1c**). In contrast, conjugation of AAV9 to AAV_DJ_ enhanced NLS-GFP expression by 13.3 times (**Fig. 1c, d**; *p*=4.10×10^−3^), demonstrating that AAV_DJ_ escorted AAV9 into target cells.

**Fig. 1.**
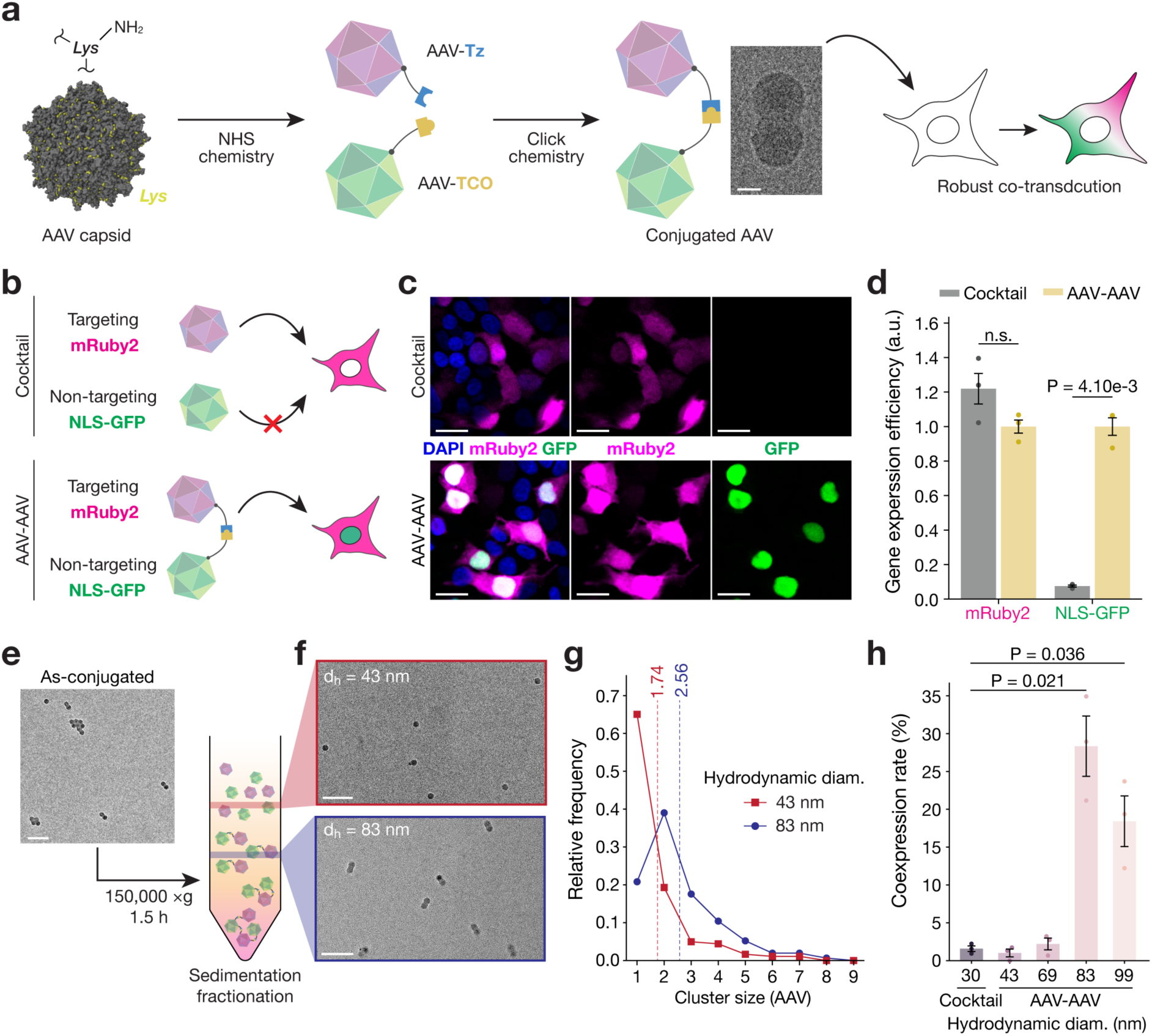
Conjugated AAVs for co-delivery of two transgenes. **a,** Scheme of the synthesis of AAV-AAV conjugates. AAVs were functionalized with either Tz or TCO through NHS chemistry, followed by conjugation through inverse electron-demand Diels-Alder click chemistry. A representative TEM image of an AAV-AAV dimer is shown. Scale bar, 10 nm. **b**, Schematics of the proof-of-concept in vitro assay with two fluorescent proteins. In the cocktail condition, two AAVs, AAV-DJ packaging pAAV-CAG-mRuby2 that transduced HEK293T cells efficiently and AAV9 packaging CAG-NLS-GFP that did not transduce HEK293T cells efficiently, were mixed and incubated with HEK293T cells. The same two AAVs were conjugated and incubated with HEK293T cells in the AAV-AAV condition. **c**, Confocal images of HEK293T cells incubated with the cocktail or AAV-AAV conjugates for 24 hours (multiplicity of infection (MOI) = 100). Blue – DAPI, Magenta – mRuby2 (AAV-DJ), Green - NLS-GFP (AAV9). Scale bars, 10 µm. **d**, mRuby2-positive and GFP-positive cell percentages in the two groups, as quantified from confocal images and with CellProfiler^57^. **e**, Schematic of sedimentation fractionation and representative TEM images of as-synthesized AAV-AAV conjugates. Scale bar, 100 nm. **f**, Representative TEM images of fractionated AAV-AAV conjugates with different hydrodynamic diameter (d_h_). Scale bars, 100 nm. **g**, Frequency of cluster sizes for the two selected fractions (d_h_ = 43, 83 nm) obtained from TEM images using a machine learning model (Detectron2; Methods). The dashed lines represent the average cluster size of each group. **h**, Percentage of cells that co-expressed mRuby2 and NLS-GFP after 24-hour incubation with the cocktail or one of the selected fractions having distinct hydrodynamic diameters, as measured by flow cytometry (MOI = 100). Bar plots are presented as mean ± standard error of the mean (S.E.M.). Statistical significance was assessed by Welch’s t-test (n = 3 biological replicates).

The physical size of individual conjugates may influence transduction efficiency, as excessively large clusters could interfere with AAV’s canonical endocytosis pathways or nuclear entry^30^. We therefore fractionated AAV_DJ_-AAV9 conjugates by size via ultracentrifugal sedimentation (**Fig. 1e**)^31^. Dynamic light scattering (DLS) confirmed separation into populations with distinct hydrodynamic diameters (d_h_; Fig. S2). Transmission electron microscopy (TEM) of selected fractions verified conjugation, and the observed cluster sizes correlated with d_h_ (**Fig. 1f, g**). Co-expression efficiency varied substantially with conjugate d_h_ (**Fig. 1h**), and the fraction exhibiting the highest co-expression rate (17.9-fold increase over the AAV cocktail; *p*=0.021) contained the greatest proportion of AAV-AAV dimers (∼40%), an architecture that ensures a 1:1 stoichiometry of the two serotypes.

We next employed AAV-AAV conjugates to deliver transgenes exceeding AAV’s packaging capacity. As a first test cargo, we selected the SpCas9 gene editing cassette (∼6.0 kb including the AAV inverted terminal repeats) and split it using split-intein domains, which mediate post-translational *trans*-splicing to reconstitute the full-length SpCas9 protein in situ (**Fig. 2a**)^32–35^. Gene editing efficiency was assessed using StopLight HEK, an established reporter cell line (Methods), in which GFP is expressed upon indel formation at the target sequence (**Fig. 2b**)^36^. Among inteins derived from diverse host proteins and organisms^37–39,23^, the *Rma* DnaE split intein exhibited the highest gene editing activity, with no significant reduction relative to wild-type SpCas9 (*p*=0.714; **Fig. 2c**, Fig. S3). The split SpCas9 constructs were separately packaged in AAV_DJ_ capsids, which were subsequently conjugated and fractionated as described above. In StopLight HEK cells at 24 hours post-transduction, AAV_DJ_-AAV_DJ_ conjugates achieved a 1.23-fold increase in gene editing efficiency over the cocktail (*p*=2.49×10^−4^; **Fig. 2d**) and required an 18.8% lower dose to reach 50% of the maximum gene editing activity (Fig. S4). Notably, the high transduction efficiency of AAV_DJ_ in HEK cells represents a stringent baseline, yet AAV_DJ_-AAV_DJ_ conjugates still yielded a 23% improvement, highlighting the benefit of vector conjugation. This enhancement generalized to a CRISPRi transcriptional repressor, dSpCas9-KRAB-MeCP2 (∼7 kb total)^40^, for which AAV_DJ_-AAV_DJ_ conjugates achieved a 1.21-fold greater gene repression than the cocktail (*p*=0.0113; Fig. S5, S6). Together, these results demonstrate that AAV-AAV conjugates enhance co-delivery of transgenes exceeding the packaging limit of individual AAV capsids.

**Fig. 2.**
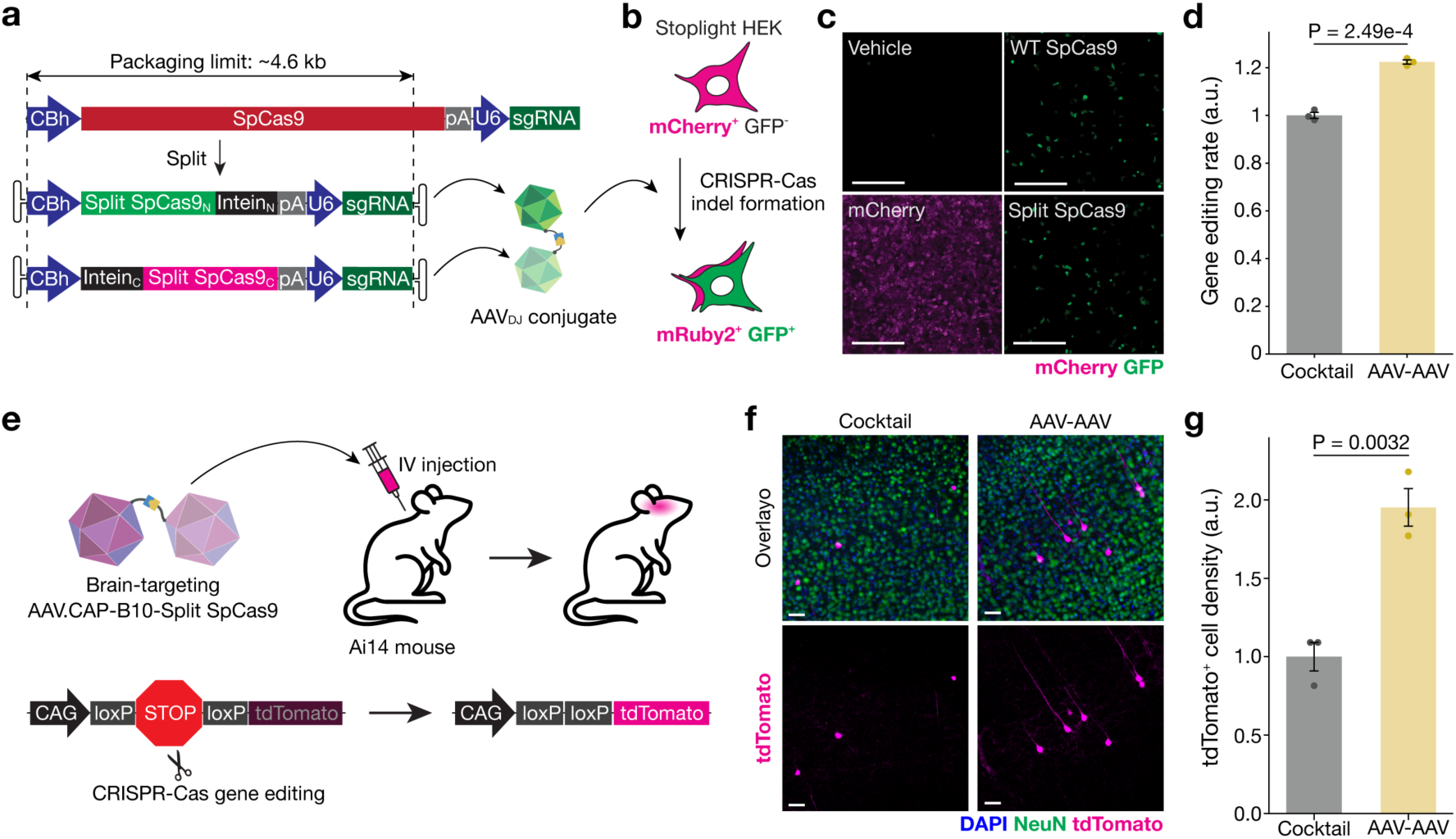
AAV-AAV conjugates for large gene delivery in vitro and in vivo. **a,** Split design of SpCas9-based gene editing cassette. The constructs were packaged separately in AAV_DJ_ capsids. **b,** StopLight reporter system employed in in vitro gene editing assays. Upon CRISPR-Cas9-mediated indel formation at the target site, GFP expression was induced by frameshift mutations. **c**, Representative confocal images of StopLight HEK cells after lipofection with wild-type SpCas9, split SpCas9 (*Rma*), or vehicle control (lipofectamine 3000). mCherry fluorescence was also shown as a reference. Scale bars, 250 µm. **d**, Gene editing efficiency of AAV_DJ_-AAV_DJ_ conjugates relative to the cocktail, as quantified by flow cytometry. **e**, Schematic of the in vivo experiment in Ai14 mice. AAV_CAP-B10_ capsids packaging the split SpCas9 constructs were conjugated and fractionated. The resulting AAV_CAP-B10_-AAV_CAP-B10_ conjugates were administered intravenously to Ai14 mice. Upon successful CRISPR-Cas9-mediated excision of the stop cassette, tdTomato expression was induced. **f**, Representative confocal images of the brain (cortex) harvested at 3 weeks post-injection from mice injected with AAV_CAP-B10_-AAV_CAP-B10_ conjugates or AAV cocktails (2.5×10^10^ vg/animal). Blue – DAPI, Green – NeuN, Magenta - tdTomato. Scale bars, 50 µm. **g**, Density of tdTomato-positive cells in the cortex, normalized to the cocktail group. Animals were administered the cocktail or AAV_CAP-B10_-AAV_CAP-B10_ conjugates (2.5×10^10^ vg/animal), and tissues were harvested at 3 weeks post-injection. Bar plots are presented as mean ± S.E.M. Statistical significance was assessed by Welch’s t-test (n = 3 biological replicates).

Building on these in vitro results, we evaluated whether AAV-AAV conjugates could enhance gene editing in vivo following systemic administration. Split SpCas9 cassettes were packaged in AAV.CAP-B10 (AAV_CAP-B10_), an AAV serotype that efficiently crosses the blood-brain barrier (BBB; **Fig. 2e**). As a readout, the Ai14 reporter mouse line was employed, in which tdTomato expression is induced upon indel formation within the stop cassette (**Fig. 2e**)^41^. A moderate AAV dose (2.5×10^10^ vg/animal) that yields extensive but unsaturated gene editing was selected to reduce systemic exposure and to reveal differences between the conjugate and cocktail conditions^7,22^. Three weeks following retro-orbital delivery, tissues were harvested, and confocal imaging was conducted on sagittal brain sections. Consistent with prior studies, tdTomato fluorescence was observed throughout the brain of mice injected with AAV_CAP-B10_ cocktail (Fig. 2f, Fig. S7). However, the expression, as quantified by the density of tdTomato-positive cells, was enhanced by 1.95 times in animals injected with AAV_CAP-B10_-AAV_CAP-B10_ conjugates as compared to the cocktail-injected controls (*p*=0.0032; **Fig. 2f-g**). This result demonstrates that AAV_CAP-B10_-AAV_CAP-B10_ conjugates enhance systemic co-delivery of split SpCas9 to the brain across the BBB.

The modularity of the CHARIOT-AAV platform motivated its extension to non-AAV vehicles for delivery of large unsplit genes. Lentiviral vectors (LVVs) integrate transgenes up to ∼10 kb into the host genome^17^, enabling stable expression throughout cell division, but do not permit cell-type-specific transduction. Building on the observation that AAV tropism can be conferred to another AAV serotype (Fig. 1b-d) and prior demonstration of AAV-mediated tissue guidance of synthetic nanomaterials^28^, we hypothesized that the targeting properties could be similarly imparted to LVVs through AAV conjugation. Unlike AAVs, LVVs are cloaked in lipid envelopes, which are not readily functionalized via carbodiimide chemistry for IEDDA reaction used in AAV-AAV conjugation. To overcome this, we employed SNAP-tag technology, in which a genetically encoded protein tag (SNAP-tag) reacts with *O*^6^-benzylguanine (BG) to form a covalent bond (**Fig. 3a**)^42^. AAV_DJ_ capsids were functionalized with BG via carbodiimide chemistry, while LVVs displaying SNAP-tag were produced by incorporating a SNAP-tag-encoding plasmid during virus packaging^25^. These LVVs, carrying a gene for GFP, also harbored mutant vesicular stomatitis virus G-protein (VSV-G) that reduced their transduction in HEK cells^25^. Following crosslinking via the SNAP-tag-BG reaction (Fig. S8), HEK293T cells were transduced with AAV_DJ_-LVV conjugates or stoichiometric vector cocktails. At 24 hours post-transduction, ∼13% of cells treated with AAV_DJ_-LVV conjugates showed GFP expression, compared with < 1% in the cocktail condition (*p*=4.24×10^−4^; **Fig. 3b, c**). A non-split SpCas9-based gene editing cassette (∼10 kb) was also delivered within AAV-LVV conjugates, which yielded a significantly higher gene editing rate than the cocktail group (Fig. S9). These data demonstrate AAV-guided delivery of a non-AAV vector carrying a large transgene.

**Fig. 3.**
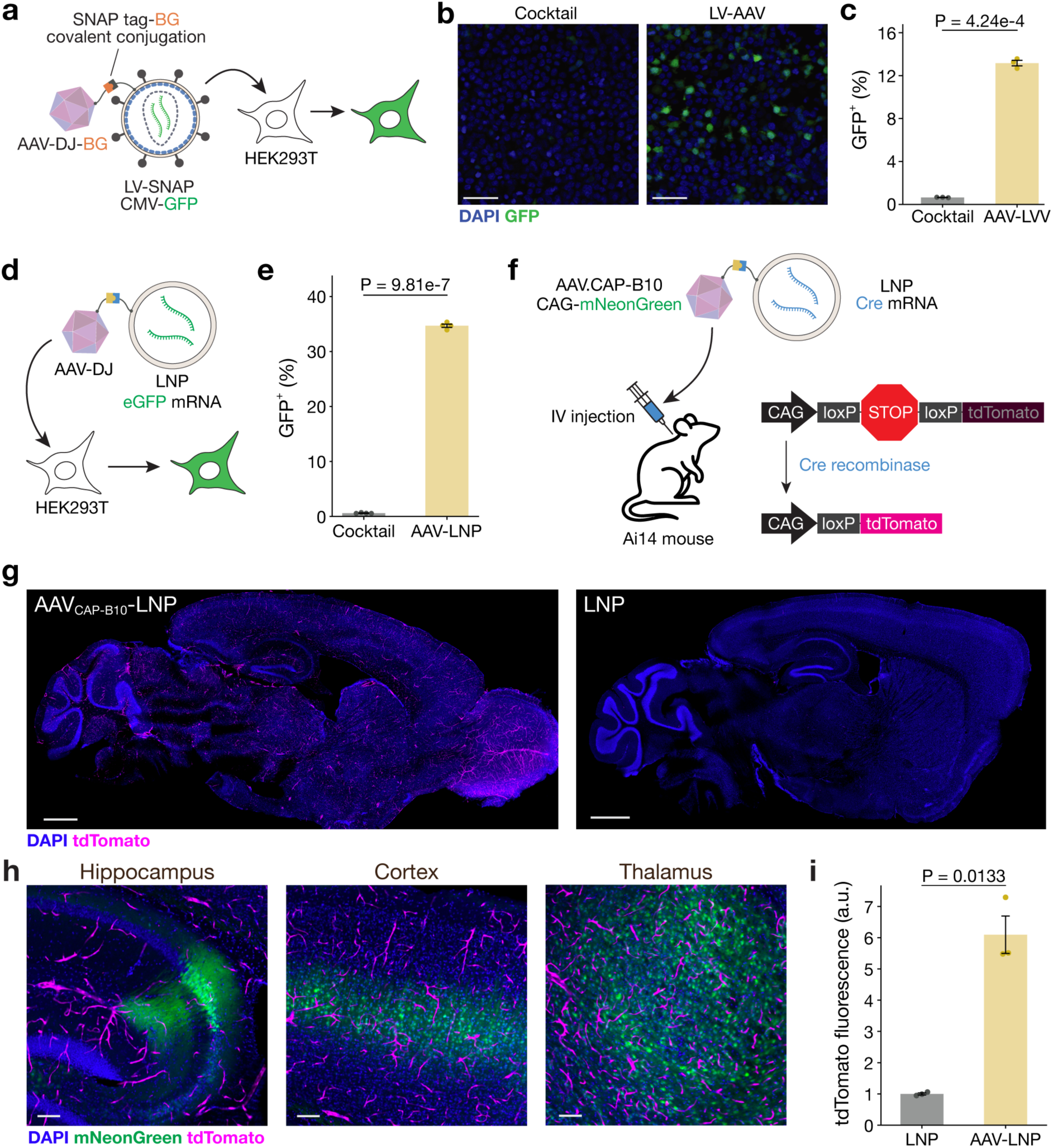
Expansion of CHARIOT-AAV to lentiviral vectors and lipid nanoparticles. **a,** Schematic of the in vitro evaluation of AAV-LVV conjugates in HEK293T cells. During LVV packaging, mutated VSV-G and SNAP-tag membrane proteins were incorporated to minimize nonspecific transduction of HEK293T cells and to render LVV surfaces reactive with BG, respectively. The LVVs packaged CMV-GFP. AAV-DJ capsids were functionalized with BG via NHS chemistry. In the cocktail group, intact AAV-DJ was used instead of BG-functionalized AAV-DJ. HEK293T cells were incubated with either LVV-AAV or LVV at a multiplicity of infection of 276. **b**, Representative confocal images of HEK293T cells transduced with a cocktail of LVVs and AAVs or AAV-LVV. HEK293T cells were fixed with 4% PFA at 24 hours post-transduction. Blue – DAPI, Green - GFP. Scale bars, 50 µm. **c**, Quantification of GFP-positive cells in the cocktail and AAV-LVV conditions. At 24 hours post-transduction, cells were harvested and analyzed by flow cytometry. **d**, Schematic of the in vitro experiment paradigm for LNP-AAV conjugates. LNPs packaged eGFP mRNA were functionalized with Tz. AAV-DJ was functionalized with NHS-PEG24-TCO. The functionalized LNP-Tz and AAV-TCO were conjugated, followed by quenching of the Tz and TCO groups. The resulting AAV-LNP conjugates were added to a culture of HEK293T cells in serum-free media to suppress ApoE-mediated transfection. At 24 hours post-transfection, HEK293T cells were harvested and analyzed for GFP fluorescence by flow cytometry. **e**, Percentage of GFP-positive cells in the cocktail and AAV-LNP groups. **f**, Schematic of brain-targeting AAV-LNP delivery in mice. LNPs packaging Cre mRNA were conjugated with AAV.CAP-B10. The AAV_CAP-B10_-LNP were administered to Ai14 mice intravenously (0.23 mg-mRNA/kg). The tissues were harvested at 5 days post-injection for confocal imaging. **g**, Confocal images of brain sagittal sections from mice injected with AAV_CAP-B10_-LNP (left) and LNP control (right). Scale bars, 1 mm. **h**, Confocal images of hippocampus, cortex, and thalamus from mice injected with AAV_CAP-B10_-LNP. Scale bars, 100 µm. **i**, tdTomato fluorescence intensity in CD31⁺ cerebral vasculature, normalized to the LNP group. Bar plots are presented as mean ± S.E.M. Statistical significance was assessed by Welch’s t-test (c, i: n = 3 biological replicates; e: n = 4 biological replicates).

Finally, we extended CHARIOT-AAV to the delivery of non-viral vectors. Lipid nanoparticles (LNPs) accommodate large genetic cargo (up to ∼10 kb) as well as molecular payloads (e.g. proteins), yet they commonly accumulate in the liver via apolipoprotein E (ApoE) adsorption^43^. Although decoration of LNPs with poly(ethylene glycol) (PEG) bearing longer alkyl chains, such as C18, prevents ApoE adsorption and hepatocyte transfection, thereby prolonging circulation time^19,44,45^, extrahepatic targeting remains limited, with only a few organs accessible to date^26,27^. We therefore sought to redirect LNPs beyond the liver using CHARIOT-AAV (**Fig. 3d**). TCO-functionalized AAVs and Tz-functionalized LNPs were conjugated via IEDDA click chemistry (Fig. S10). DLS measurements showed a slight increase in hydrodynamic diameter from 101.6 nm (cocktail) to 108.8 nm (AAV-LNP) upon conjugation (Fig. S11), confirming that excessive crosslinking did not occur. GFP mRNA was packaged in LNPs, and GFP-positive cells were quantified by flow cytometry at 24 hours post-transfection. To minimize ApoE adsorption and suppress AAV-independent transfection, serum-free medium was used during the procedure. In the AAV_DJ_-LNP conjugate group, 43.3 ± 4.5% of HEK cells expressed GFP, compared with 0.6 ± 0.1% for the cocktail group (**Fig. 3e**), demonstrating that covalent AAV conjugation enables targeted cellular uptake of LNPs.

Given their favorable long-term safety profile^46^, LNPs are a promising vehicle for delivery of molecular therapeutics in vivo. To evaluate the ability of CHARIOT-AAV to guide LNPs toward the brain, we conjugated AAV_CAP-B10_ to LNPs formulated with C18-PEG lipids, which suppress unwanted hepatic uptake (**Fig. 3f**). In these conjugates, AAV_CAP-B10_ capsids packaged a fluorescent protein mNeonGreen gene driven by a CAG promoter, and LNPs harbored Cre recombinase mRNA. The ability of AAV_CAP-B10_-LNP conjugates to deliver molecular payloads to target tissues was then evaluated in Ai14 mice, which carry a Cre-dependent tdTomato reporter allele driven by the ubiquitous CAG promoter. AAV_CAP-B10_-LNP conjugates and unfunctionalized LNPs were administered intravenously, and tissues were harvested five days post-injection. In the brain, tdTomato expression localized to the vasculature, with significantly higher fluorescence intensity in CD31⁺ cerebral vessels compared to LNP-only controls (p=0.0133; **Fig. 3g-i**, Fig. S12), suggesting that endothelial internalization was driven by the conjugated AAV_CAP-B10_ capsids. Notably, the mNeonGreen carried by the AAV_CAP-B10_ was expressed exclusively in the brain parenchyma rather than endothelial cells (**Fig. 3h**). As expected, tdTomato expression was also observed in the liver, consistent with residual hepatic uptake of LNPs despite C18-PEG incorporation (Fig. S13). Importantly, mNeonGreen expression in the liver was minimal, suggesting that AAV_CAP-B10_-LNP conjugates were preferentially directed toward the brain rather than the liver. These data demonstrate that AAV capsid conjugation redirects non-AAV vehicles toward defined cell populations in a serotype-dependent manner both in vitro and in vivo.

Here we introduced CHARIOT-AAV, a tissue-specific systemic delivery platform for genetic payloads beyond AAV’s packaging capacity. AAV-AAV conjugates achieved systemic brain transduction with split SpCas9 across the BBB. This design is readily adaptable to other dual-AAV systems, thereby providing a generalizable platform for large gene delivery. AAV-LVV and AAV-LNP conjugates extend this capability to unsplit full-length genes up to ∼10 kb. AAV-LNP further enables transient expression of mRNAs and the delivery of molecular payloads (e.g. proteins), which are advantageous for gene-therapy applications to minimize off-target effects. For even larger genes (e.g. full-length dystrophin cDNA for Duchenne muscular dystrophy therapy, >11 kb), conjugation of three or more AAVs holds promise. In preliminary studies, co-delivery of three distinct fluorescent proteins was achieved in HEK293T cells using three-AAV conjugates (Fig. S14), and the *Rma* DnaE and *Npu* DnaE inteins showed no cross-reactivity (Fig. S3), enabling orthogonal splitting of a large gene across three vectors.

Following intravenous injection, AAV_CAP-B10_-LNP conjugates achieved spatially distinct delivery: LNP payloads to brain endothelial cells and AAV payloads to the parenchyma. This dual targeting holds therapeutic implications, as brain endothelial cells are a major cellular component of the BBB, which is disrupted in several neurodegenerative^47–49^ and neurodevelopmental^50^ conditions. These data also provide mechanistic insight into the fate of LNPs during AAV_CAP-B10_-mediated transcytosis. The conjugated LNPs likely release their mRNA cargo within endothelial cells via the LNPs’ intrinsic endosomal escape machinery. To extend LNP payload delivery beyond the endothelium, future efforts could engineer LNPs with delayed endosomal escape kinetics or employ alternative vector systems that lack active endosomal escape mechanisms. While scalable manufacturing, including stoichiometry control, remains to be established for therapeutic applications, CHARIOT-AAV reconceptualizes AAV capsids as modular targeting elements. By decoupling tissue-specific tropism from cargo capacity, this platform enables tissue- and cell-type-specific delivery of diverse therapeutics and research tools beyond the limitations of individual vectors.

## Methods

### Reagents

Sucrose (BioXtra grade, ≥99.5%; S7903) was purchased from Sigma-Aldrich. *O*^6^-benzylguanine (BG)-GLA-NHS (#S9151S) and BG-PEG-NH_2_ (#S9150S) were purchased from New England Biolabs. Methoxy poly(ethylene glycol)24-NHS (mPEG24-NHS; #BP-23970), *trans*-cycrooctene-PEG24-NHS (TCO-PEG24-NHS; #BP-26353), tetrazine-PEG5-NHS ester (Tz-PEG5-NHS; #BP22681), mPEG-methyltetrazine (Mw 2000; mPEG-Tz; #BP-26352), and mPEG4-TCO (#BP27872) were purchased from BroadPharm. All lipid components used for lipid nanoparticle (LNP) formulation were purchased from commercial suppliers. The ionizable lipid ALC-0315 (#890900), 1,2-distearoyl-sn-glycero-3-phosphocholine (DSPC; #850365), 1,2-dimyristoyl-sn-glycero-3-phosphoethanolamine-N-[methoxy(polyethylene glycol)-2000] (DMG-mPEG2000; #880151), and 1,2-distearoyl-sn-glycero-3-phosphoethanolamine-N-[methoxy(polyethylene glycol)-2000] (DSG-mPEG2000; #880152) were obtained from Avanti Polar Lipids. Cholesterol was purchased from Millipore Sigma (#C8667). The functionalized PEG-lipid 1,2-distearoyl-sn-glycero-3-phosphoethanolamine-N-[methoxy(polyethylene glycol)-2000]-methyltetrazine was purchased from BroadPharm (DSPE-PEG2000-methyltetrazine; #BP-43749). N1-Methylpseudouridine (N1mU)-modified mRNAs encoding Cre recombinase and enhanced green fluorescent protein (eGFP) were purchased from TriLink Biotechnologies (#L-8111) and GenScript, respectively (#RP-A00009).

### Cloning of constructs

NEB 5-alpha Competent E. coli (High Efficiency; NEB C2987H) was used in transformation for all genetic constructs. Miniprep was performed using NucleoSpin Plasmid EasyPure (Takara Bio #740727.50). All plasmid sequences were confirmed by whole-plasmid sequencing performed by Plasmidsaurus. Parent plasmids are listed in Table 1, and new construct employed in this study can be found in the Supplementary Information.

**Table 1.** Base plasmids.

| Name | Source | Comment |
| --- | --- | --- |
| pUCmini-iCAP-AAV.CAP-B10 | Addgene plasmid #175004 | Gifts from Viviana Gradinaru |
| pAAV-CAG-mRuby2 | Addgene plasmid #99123 | Gifts from Viviana Gradinaru |
| pAAV-CAG-NLS-GFP | Addgene plasmid #104061 | Gifts from Viviana Gradinaru |
| pAAV-CAG-mTurquoise2 | Addgene Plasmid #99122 | Gifts from Viviana Gradinaru |
| pAAV2/9n | Addgene plasmid #112865 | Gift from James M. Wilson |
| pMK231 (AAVS1 CMV-MCS-PURO) | Addgene plasmid #105924 | Gift from Masato Kanemaki |
| PX458-AAVS1 | Addgene plasmid #113194 | Gift from Adam Karpf |
| AAVS1-Pur-CAG-EGFP | Addgene plasmid #80945 | Gift from Su-Chun Zhang |
| N-Terminal Split Cas9 with GyrA intein | Addgene plasmid #58693 | Gift from Gang Bao |
| C-Terminal Split Cas9 with GyrA intein | Addgene plasmid #58694 | Gift from Gang Bao |
| dCas9-KRAB-MeCP2 | Addgene plasmid #110821 | Gift from Alejandro Chavez & George Church |
| pX330-U6-Chimeric_BB-CBh-hSpCas9 | Addgene plasmid # 42230 | Gift from Feng Zhang |
| pHCMV-VSVgSS-SNAP-TM | Addgene plasmid #207319 | Gift from Feng Zhang |
| pMD2.G-K47Q/R354Q | Addgene plasmid #207323 | Gift from Feng Zhang |
| **lentiCRISPR v2** | Addgene plasmid # 52961 | Gift from Feng Zhang |
| pMDLg/pRRE | Addgene plasmid #12251 | Gift from Didier Trono |
| pRSV-Rev | Addgene plasmid #12253 | Gift from Didier Trono |
| pLenti CMV GFP Puro (658-5) | Addgene plasmid #17448 | Gift from Eric Campeau & Paul Kaufman |
| pAAV-DJ | Cell Biolabs, Inc. #VPK-420-DJ |  |
| pHelper | Cell Biolabs, Inc. |  |
| RV01-pRGS-Reporter with GFP | PNA bio |  |

To construct StopLight HEK, synthetic gene fragments containing mCherry-F2A-stop-EGFP-stop-EGFP (IDT, gBlocks Gene Fragments)^36^ were inserted into pMK231 (AAVS1-CMV-MCS-PURO, Addgene #105924) via Gibson assembly using NEBuilder HiFi DNA Assembly Master Mix (NEB #E2621L).

To create split SpCas9, the *Rma*^51^, *Npu*^38,52^, and *Cfa*^39^ DnaE split intein genes were synthesized (IDT, gBlocks Gene Fragments). Split SpCas9 sequences were amplified from pX330, and these fragments were assembled into the pAAV backbone including an sgRNA cassette (see Supplementary Methods).

The split dSpCas9-KRAB-MeCP2 constructs were generated from dCas9-KRAB-MeCP2 (Addgene #110821). *Rma* DnaE split intein sequences were amplified from the split SpCas9 (*Rma*) constructs by PCR using Q5 Hot Start High-Fidelity 2X Master Mix (NEB #M0494S). dCas9-KRAB-MeCP2 was also split at several sites (L833/S834, Q844/S845, D850/S851, R859/S860, K867/S868, P872/S873, and L909/S910) by PCR. The resulting gene fragments were assembled via Gibson assembly (3 fragments for the C-terminal split construct; 5 fragments for the N-terminal split construct; see Supplementary Methods).

The sgRNA sequence for the StopLight HEK experiment was obtained from the original article^36^. The same sgRNA sequence was used for both split constructs. The sgRNA sequences for the CAG promoter were designed using CRISPick^53,54^, and the top two sequences were used in the two split constructs. The sgRNA sequence for Ai14 mouse experiments was obtained from a previous study^41^.

### Cell culture and cloning of GFP-expressing and StopLight HEK293T

HEK293T cells were cultured in Dulbecco’s Modified Eagle Medium (DMEM, Gibco #10569044) with 10% fetal bovine serum (FBS, Cytovia #SH30396.03HI). 1% Penicillin-Streptomycin (P/S) was also supplemented except for lentiviral vector production. Cells were passaged at 90% confluency with TrypLE Express (Gibco #12605028).

To obtain GFP-expressing HEK and Stoplight HEK cells, HEK293T cells were cultured in 6-well plates in DMEM with 10% FBS and 1% P/S and dissociated with TrypLE Express for 5 min. Then, a total of 2×10^6^ cells were co-transfected with 10 µg of PX458-AAVS1 plasmid, which expresses SpCas9, and 6 µg of AAVS1-CMV-Stoplight-Puro or AAVS1-Pur-CAG-EGFP in 100 µL of Opti-MEM (Gibco #31985088). Electroporation was performed using a BTX ECM830 electroporator (Poring plus: voltage, 150 V; pulse length, 5 ms; pulse, 100 ms; number of pulses, 2. Transfer pulse: voltage, 20 V; pulse length, 100 ms; pulse, 100 ms; number of pulses, 5). After electroporation, HEK293T cells were seeded in DMEM with 10% FBS and 1% P/S, and 24 hours post-electroporation, transfected cells were treated with puromycin at 1.5 μg/ml to select genome-edited cells. Then, the cells were further purified with a cell sorter (FACS BD Melody) according to mCherry or EGFP positive expression, followed by genotyping to validate the correct genome insertion and sequence.

### Evaluation of split constructs by lipofection

Prepared split constructs were evaluated in StopLight HEK or GFP-expressing HEK cells by lipofection. Wild-type constructs (i.e. SpCas9, dSpCas9-KRAB-MeCP2) were used as benchmarks. Cells were prepared in a 96-well plate and used at ∼80% confluency. 6 µL of Lipofectamine 3000 (Thermo Fisher Scientific #L3000001) and 150 µL of Opti-MEM (Gibco #31985062) was mixed. In another tube, 150 µL of Opti-MEM, 6 µL of P3000 reagent, and 75 fmol of plasmid DNAs were mixed. The two solutions were mixed and incubated at room temperature for 10 min, and then added to 3 wells (100 µL/well). At 48 hours post-lipofection, the cells were harvested and analyzed by flow cytometry (BD FACSymphony A3).

### AAV packaging

AAVs were packaged in the lab following an established protocol^55^. Briefly, HEK293T cells were cultured in DMEM with 10% FBS and 1% P/S. At 95% confluency, triple transfection with a capsid plasmid, a helper plasmid, and a plasmid encoding the gene of interest within the pAAV backbone was performed using PEI Max. The medium was exchanged with fresh DMEM with 5% FBS at 10 hours post-transfection. The culture medium was harvested at 72 hours post-transfection and fresh 5% FBS DMEM was added. At 120 hours post-transfection, both medium and cells were harvested. AAV capsids inside the cells were released using salt active nuclease (HL-SAN, ArcticZyme #70910-202). AAV capsids in the medium were collected by PEG-mediated precipitation. Collected AAV capsids were purified by ultracentrifugation (350,000 ×g, 2 hours, 25 min, 18 °C) in an iodixanol density gradient (OptiPrep, STEMCELL TECHNOLOGIES #07820). Purified AAVs were stored in Dulbecco’s phosphate-buffered saline (DPBS) (Corning #20-031-CV) with 0.1% Pluronic F-68 (Gibco #24040032) (DPBS-F68). AAV titers were measured by quantitative PCR (qPCR) using an AAV real-time PCR titration kit (Takara Bio #6233).

### AAV modification

AAV capsids were functionalized with Tz, TCO, or BG, depending on the CHARIOT-AAV system, via NHS chemistry following a previously reported protocol^28^. Briefly, AAVs were suspended in pH 8.4 sodium bicarbonate solution in DPBS-F68 using filter centrifuge units (Amicon Ultra Centrifugal Filter, 100 kDa MWCO, Millipore #UFC510024). 2 pmol of AAV capsids were reacted with ligands at a ligand-to-lysine residue ratio of 0.1 (AAV-DJ and AAV9) and 0.075 (AAV.CAP-B10). The solution was allowed to react at 4°C overnight. The resulting functionalized AAVs were purified with DPBS-F68 using the filter centrifuge units. Once purified, functionalized AAVs were titrated by qPCR.

### AAV-AAV conjugation and fractionation

Tz-functionalized AAV capsids and TCO-functionalized AAV capsids were reacted in DPBS-F68 and conjugated via the inverse electron-demand Diels-Alder cycloaddition click chemistry. The ratio of the two capsids was fixed at 1:1. Typically, functionalized AAV solutions were adjusted to 10 nM; 50 µL of each AAV solution was then mixed so that the final concentration of each AAV was 5 nM. The solution was stored at 4 °C overnight. To stop the reaction, 50 µL of mPEG4-TCO (0.1 mg/mL) was first added and allowed to react for 4 hours at 4 °C. Subsequently, 140 µL of mPEG2k-Tz (1 mg/mL) was added and allowed to react for another 4 hours at 4 °C. The resulting AAV-AAV conjugates were purified by medium exchange with DPBS-F68 using a filter centrifuge unit (Amicon Ultra Centrifugal Filter, 0.5 mL, 100 kDa MWCO, Millipore #UFC510024). The volume was adjusted to ∼200 µL after the final centrifugation step, then loaded onto a sucrose gradient (8%-4 mL, 10%-4 mL, 12.5%-6 mL, 15%-6 mL, 17.8%-4 mL, 20%-2 mL, 25%-2 mL, and 40%-2 mL; in a 30 mL sterile, open-top ultracentrifuge tube (Beckman Coulter # C14307)). The solution was centrifuged in an SW 32 Ti Swinging-Bucket Rotor (Beckman Coulter) at 150,000 ×g for 1.5 hours at 4 °C. The solution was fractionated into 1 mL from the bottom of the tube using a syringe, and the titer of each fraction was determined by qPCR. Prior to DLS measurement, the fractionated solutions were medium-exchanged with DPBS-F68 to remove remaining sucrose. The titer was again determined after the medium exchange step.

Transmission electron microscopy of AAV-AAV conjugates was performed on an FEI G2 Spirit TWIN TEM. Fifty images were collected for each sample. Conjugate size analysis was performed using Detectron2 (model: Mask R-CNN X101-FPN, model ID: 139653917)^56^. The model was trained with human-annotated data (TEM images of AAV particles, ∼1000 particles in total).

### AAV-AAV transduction test in vitro and in vivo

For in vitro experiments, fractionated AAV-AAV conjugates were added to a culture of HEK293T, StopLight HEK, or GFP-expressing HEK cells at varying MOI values. Transduction durations were 24 hours for the two fluorescent protein transduction tests (Fig. 1) and the split SpCas9 gene editing tests (Fig. 2) and 72 hours for the CAG promoter suppression test (Fig. S6). Cells were harvested and measured by flow cytometry (BD FACSymphony A3).

For in vivo experiments, AAV-AAV conjugates were intravenously injected via the retro-orbital route into 6-week-old Ai14 mice (2 males and 1 female; 2.5×10^10^ vg/animal). Three weeks post injection, the animals were euthanized and perfused with PBS and 4% paraformaldehyde (PFA) in PBS. Harvested tissues were further fixed in 4% PFA overnight. Fixed tissues were sectioned at a thickness of 50 µm in the sagittal plane using a Vibratome. The obtained brain slices were permeabilized in 0.3% Triton X-100 for 30 min, followed by blocking with 3% normal donkey serum (NDS) for 2 hours at room tempearture. Primary antibody labeling was performed in 3% NDS at 4 °C overnight (Anti NeuN antibody (host: rabbit), 1:500, Proteintech #26975-1-AP). After three cycles of washing with PBS, the slices were stained with secondary antibodies in 3% NDS for 2 hours at room temperature (Donkey anti-Rabbit IgG (H+L) Highly Cross-Adsorbed Secondary Antibody, Alexa Fluor 647, 1:1,000, Thermo Fisher Scientific #A-31573). The sections were subsequently stained with DAPI for 30 min (1:20,000), and then mounted on glass slides using Fluoromount-G. Confocal imaging was performed on Leica Stellaris 5 with a 10x objective lens. CellProfiler^57^ was used for quantification.

For both in vitro and in vivo experiments, intact AAVs were used for the cocktail conditions, and the solutions were adjusted to the same titer before use.

### Lentiviral vector production

HEK293T cells were prepared in DMEM with 10% FBS in three 15-cm plates and were transfected with a mixture of plasmids using PEI Max (Polysciences, #24765). Packaging plasmids (pMDLg/pRRE (Addgene 12251) and pRSV.Rev (Addgene 12253)), a transfer plasmid (pLenti CMV GFP Puro (658-5) (Addgene 17448)), and envelope plasmids (pMD2.G-K47Q/R354Q (Addgene 207323) and pHCMV-VSVgSS-SNAP-TM (Plasmid 207319)) were mixed at the ratio of pMDLg/pRRE:pRSV.Rev:transfer plasmid:pMD2.G-K47Q-R354Q:pHCMV-VSVgSS-SNAP-TM=1:1:3:0.75:0.25. At 8 hours post-transfection, the media was exchanged with fresh DMEM with 10% FBS. At 48 hours post-transfection, the media containing lentiviral vectors (LVVs) was harvested, followed by filtration with a 0.45 µm PES membrane filter. The solution was then concentrated by ultracentrifugation (50,000 ×g, 30 min, 4 °C). The viral pellet was resuspended in PBS. The titer of LVVs was determined by RT-qPCR (Lenti-X qRT-PVR Titration Kit; #631235, Takara bio).

### AAV-LVV conjugation and in vitro transduction test

SNAP-tag-functionalized LVVs and BG-functionalized AAVs were conjugated in DPBS-F68 at an AAV/LVV ratio of 100 (100 AAVs per LLV). The reaction was stopped at different time points by adding mPEG24-BG, which was prepared by mixing BG-PEG-NH2 and mPEG24-NHS at a molar ratio of 1:1.5 in pH 8.4 sodium bicarbonate in DPBS-F68 at 4 °C overnight. Based on the optimization results (Fig. S9), 4 hours was selected as the reaction time. Prepared AAV-LVV conjugates were tested in HEK293T or StopLight HEK cells at an MOI of 150. 24 hours post-transduction, HEK cells were harvested and analyzed for GFP fluorescence by flow cytometry (BD FACSymphony A3).

### Lipid nanoparticle production

Lipid nanoparticles (LNPs) were prepared via microfluidic mixing as previously described^58^. Briefly, an ethanol phase containing ALC-0315, DSPC, cholesterol, PEG-lipid, and functionalized PEG-lipid (DSPE-PEG2000-mTztetrazine) was prepared. The molar ratio of ionizable lipid, DSPC, cholesterol, and total PEG-lipid (including functionalized PEG-lipid) was 50:10:38.5:1.5. For *in vitro* experiments, the PEG-lipid component consisted of DMG-mPEG2000, whereas DSG-mPEG2000 was used for *in vivo* studies. The DSPE-PEG2000-mTz was included at a concentration of 0.005-0.5 mol% of the total 1.5 mol% PEG-lipid content.

This ethanol phase was rapidly mixed with an aqueous solution of RNA dissolved in 10 mM citrate buffer (pH 3.0) at a volumetric aqueous-to-ethanol ratio of 3:1. Following microfluidics mixing, LNPs were dialyzed for at least 18 hours at 4 °C against phosphate-buffered saline (PBS) using Slide-A-Lyzer dialysis devices with a 10 kDa molecular weight cut-off (ThermoFisher). Nanoparticle size and polydispersity index (PDI) were measured using a Malvern Zetasizer. RNA encapsulation efficiency was assessed via the Quant-iT RiboGreen assay (ThermoFisher).

### AAV-LNP conjugation and transduction evaluation

Tz-functionalized LNPs and TCO-functionalized AAVs were conjugated through the IEDDA click chemistry in DPBS-F68. The number ratio of LNP particles and AAV capsids was 1:0.75 (LNP:AAV), and the concentrations of LNPs and AAVs were 20 nM and 10 nM, respectively. The reaction was quenched at different time points by adding Tz-PEG2k and mPEG4-TCO subsequently. Typically, 100 µL of LNP solution and 150 µL of AAV solution was mixed and allowed to react for 15 min at 4 °C. The functional groups were quenched with 150 µL of mPEG2k-Tz (0.01 mg/mL) for 4 hours at 4 °C and subsequently with 270 µL of mPEG4-TCO (0.1 mg/mL) for 4 hours at 4 °C. The resulting AAV-LNP conjugates were dialyzed for overnight at 4 °C against DPBS-F68 using Slide-A-Lyzer dialysis devices with a 100 kDa molecular weight cut-off (ThermoFisher).

In in vitro experiments, HEK293T cells were treated with AAV-LNP conjugates in serum-free DMEM. The LNP particle-to-cell ratio was 3160. Two hours post-transfection, the medium was exchanged with 10% FBS DMEM. At 24 hours post-transfection, cells were harvested and measured for GFP expression by flow cytometry (BD FACSymphony A3).

In in vivo experiments, AAV-LNP conjugates were injected into Ai14 mice intravenously via the retro-orbital route (2 males and 1 female per group; 0.23 mg mRNA/animal). Five days post-injections, the mice were euthanized and perfused with PBS and 4% PFA. The harvested tissues were further fixed in 4% PFA overnight. The brain tissues were sectioned in a sagittal plane at 50 µm using a Vibratome. The brain slices were permeabilized with 0.3% Triton X-100 for 30 min, followed by blocking with 3% normal donkey serum (NDS) at 4 °C overnight. The tissues were then labeled with anti-CD31 antibody (Rat Anti-Mouse CD31, BD Pharmingen 550274, 1:200) at 4 °C overnight. Secondary antibody staining was performed at room temperature on a shaker for 1 hour (Donkey anti-Rat IgG (H+L) Highly Cross-Adsorbed Secondary Antibody, Alexa Fluor Plus 647, Thermo Fisher Scientific A48272, 1:1000). After 2 washing cycles with PBS, the brain slices were incubated with DAPI (1:20,000) for 3 min. Following 2 washing cycles, the sections were mounted on glass slides using Fluoromount-G. Confocal imaging was performed on Leica Stellaris 5 with a 10x objective lens. Quantification of fluorescence was conducted using CellProfiler4^57^.

## Acknowledgement

We thank the Koch Institute’s Robert A. Swanson (1969) Biotechnology Center for technical support, C. Hallee (the Genomics Core) and the Flow Cytometry Core. This work made use of the MRSEC Shared Experimental Facilities at MIT, supported by the National Science Foundation under award number DMR-1419807. This work was funded in part by the Pioneer Award from the National Institutes of Health and National Institute for Complementary and Integrative Health (DP1-AT011991, P.A.), McGovern Institute for Brain Research at MIT, K. Lisa Yang and Hock E. Tan Center for Molecular Therapeutics at MIT (P.A.), NIH grants R01MH085802 and R01NS130361 (M.S.), MURI grant W911NF2110328 (M.S.), the Picower Institute Innovation Fund (M.S.), and the Simons Foundation Autism Research Initiative through the Simons Center for the Social Brain (M.S.). R.J.M. acknowledges support from the Army Research Office under award W911NF-23-2-0101. K.N. is a recipient of a scholarship from the Honjo International Scholarship Foundation and the Y. Eva Tan Fellowship Program. J.L.B. acknowledges support by the Schmidt Science Fellows program, in partnership with the Rhodes Trust. S.T., C.G.L., and D.G.A. acknowledge support from the sponsored research funding from FUJIFILM Corporation.

## Author Contributions

K.N., T.O. and P.A. designed the study. K.N. and E.V.P. produced AAVs and LVVs. K.N. prepared and characterized AAV-AAV, AAV-LVV, and AAV-LNP conjugates and performed in vivo experiments. K.N. and T.O. performed in vitro experiments and cloning. T.O. prepared StopLight HEK and GFP-expressing HEK cells. S.T. and C.G.L. prepared and characterized LNPs. K.N. and E.C.G. trained the machine learning model for TEM image analysis. S.M. prepared Ai14 mice for in vivo experiments. K.N., E.V.P., and J.L.B. prepared cells in vitro experiments. K.N., T.O., M.S., D.G.A., and P.A. analyzed the data. All authors have contributed to the writing of the manuscript.

## References

1. Madigan, V., Zhang, F. & Dahlman, J. E. Drug delivery systems for CRISPR-based genome editors. Nat Rev Drug Discov 22, 875–894 (2023).

2. Mitchell, M. J. et al. Engineering precision nanoparticles for drug delivery. Nat Rev Drug Discov 20, 101–124 (2021).

3. Wang, D., Tai, P. W. L. & Gao, G. Adeno-associated virus vector as a platform for gene therapy delivery. Nat Rev Drug Discov 18, 358–378 (2019).

4. Li, C. & Samulski, R. J. Engineering adeno-associated virus vectors for gene therapy. Nat Rev Genet 21, 255–272 (2020).

5. Sahel, J.-A. et al. Partial recovery of visual function in a blind patient after optogenetic therapy. Nat Med 27, 1223–1229 (2021).

6. Wang, J.-H., Gessler, D. J., Zhan, W., Gallagher, T. L. & Gao, G. Adeno-associated virus as a delivery vector for gene therapy of human diseases. Sig Transduct Target Ther 9, 78 (2024).

7. Goertsen, D. et al. AAV capsid variants with brain-wide transgene expression and decreased liver targeting after intravenous delivery in mouse and marmoset. Nat Neurosci (2021) doi:10.1038/s41593-021-00969-4.

8. Tabebordbar, M. et al. Directed evolution of a family of AAV capsid variants enabling potent muscle-directed gene delivery across species. Cell 184, 4919–4938.e22 (2021).

9. Moyer, T. C. et al. Highly conserved brain vascular receptor ALPL mediates transport of engineered AAV vectors across the blood-brain barrier. Molecular Therapy 33, 3902–3916 (2025).

10. Chuapoco, M. R. et al. Adeno-associated viral vectors for functional intravenous gene transfer throughout the non-human primate brain. Nat. Nanotechnol. 18, 1241–1251 (2023).

11. Chen, X. et al. Engineered AAVs for non-invasive gene delivery to rodent and non-human primate nervous systems. Neuron 110, 2242–2257.e6 (2022).

12. Wu, Z., Yang, H. & Colosi, P. Effect of Genome Size on AAV Vector Packaging. Molecular Therapy 18, 80–86 (2010).

13. Xu, X. et al. Engineered miniature CRISPR-Cas system for mammalian genome regulation and editing. Molecular Cell 81, 4333–4345.e4 (2021).

14. Rauch, B. J. et al. Single-AAV CRISPR editing of skeletal muscle in non-human primates with NanoCas, an ultracompact nuclease. Preprint at 10.1101/2025.01.29.635576 (2025).

15. Thelen, J. L. et al. Morphological Characterization of Self-Amplifying mRNA Lipid Nanoparticles. ACS Nano 18, 1464–1476 (2024).

16. Geall, A. J. et al. Nonviral delivery of self-amplifying RNA vaccines. Proc. Natl. Acad. Sci. U.S.A. 109, 14604–14609 (2012).

17. Kumar, M., Keller, B., Makalou, N. & Sutton, R. E. Systematic Determination of the Packaging Limit of Lentiviral Vectors. Human Gene Therapy 12, 1893–1905 (2001).

18. Bouard, D., Alazard-Dany, N. & Cosset, F. Viral vectors: from virology to transgene expression. British J Pharmacology 157, 153–165 (2009).

19. Hou, X., Zaks, T., Langer, R. & Dong, Y. Lipid nanoparticles for mRNA delivery. Nat Rev Mater 6, 1078–1094 (2021).

20. Mittas, D. M. et al. Dual AAV vectors for efficient delivery of large transgenes. Nat Protoc (2025) doi:10.1038/s41596-025-01243-8.

21. Lin, J. et al. AAVLINK: A potent DNA-recombination method for large cargo delivery in gene therapy. Cell (2026) doi:10.1016/j.cell.2025.12.039.

22. Davis, J. R., et al. Efficient in vivo base editing via single adeno-associated viruses with size-optimized genomes encoding compact adenine base editors. Nat. Biomed. Eng 6, 1272–1283 (2022).

23. Chen, Y., et al. Development of Highly Efficient Dual-AAV Split Adenosine Base Editor for In Vivo Gene Therapy. Small Methods 4, 2000309 (2020).

24. Kuzmin, D. A. et al. The clinical landscape for AAV gene therapies. Nat Rev Drug Discov 20, 173–174 (2021).

25. Strebinger, D. et al. Cell type-specific delivery by modular envelope design. Nat Commun 14, 5141 (2023).

26. Cheng, Q. et al. Selective organ targeting (SORT) nanoparticles for tissue-specific mRNA delivery and CRISPR–Cas gene editing. Nat. Nanotechnol. 15, 313–320 (2020).

27. Chen, M. Z. et al. A versatile antibody capture system drives specific in vivo delivery of mRNA-loaded lipid nanoparticles. Nat. Nanotechnol. 20, 1273–1284 (2025).

28. Nagao, K., Macfarlane, R. J. & Anikeeva, P. Adeno-associated viruses escort nanomaterials to specific cells and tissues. bioRxiv (2025) doi: 10.1101/2025.04.04.647267.

29. Grimm, D. et al. In Vitro and In Vivo Gene Therapy Vector Evolution via Multispecies Interbreeding and Retargeting of Adeno-Associated Viruses. J Virol 82, 5887–5911 (2008).

30. Burns, A. & Datta, S. Improving Aggregation Control of Recombinant Adeno-Associated Virus Serotype 2 (rAAV2) With Small Sugars and Ionic Salts. Biotechnology Journal 20, e70157 (2025).

31. Ayuso, E. et al. High AAV vector purity results in serotype- and tissue-independent enhancement of transduction efficiency. Gene Ther 17, 503–510 (2010).

32. Shah, N. H. & Muir, T. W. Inteins: nature’s gift to protein chemists. Chem. Sci. 5, 446–461 (2014).

33. Wood, D. W., Belfort, M. & Lennon, C. W. Inteins—mechanism of protein splicing, emerging regulatory roles, and applications in protein engineering. Front. Microbiol. 14, 1305848 (2023).

34. Wang, H., Wang, L., Zhong, B. & Dai, Z. Protein Splicing of Inteins: A Powerful Tool in Synthetic Biology. Front. Bioeng. Biotechnol. 10, 810180 (2022).

35. Tasfaout, H. et al. Split intein-mediated protein trans-splicing to express large dystrophins. Nature 632, 192–200 (2024).

36. De Jong, O. G. et al. A CRISPR-Cas9-based reporter system for single-cell detection of extracellular vesicle-mediated functional transfer of RNA. Nat Commun 11, 1113 (2020).

37. Fine, E. J. et al. Trans-spliced Cas9 allows cleavage of HBB and CCR5 genes in human cells using compact expression cassettes. Sci Rep 5, 10777 (2015).

38. Truong, D.-J. J. et al. Development of an intein-mediated split–Cas9 system for gene therapy. Nucleic Acids Res 43, 6450–6458 (2015).

39. Stevens, A. J. et al. Design of a Split Intein with Exceptional Protein Splicing Activity. J. Am. Chem. Soc. 138, 2162–2165 (2016).

40. Yeo, N. C. et al. An enhanced CRISPR repressor for targeted mammalian gene regulation. Nat Methods 15, 611–616 (2018).

41. Staahl, B. T. et al. Efficient genome editing in the mouse brain by local delivery of engineered Cas9 ribonucleoprotein complexes. Nat Biotechnol 35, 431–434 (2017).

42. Juillerat, A. et al. Directed Evolution of O6-Alkylguanine-DNA Alkyltransferase for Efficient Labeling of Fusion Proteins with Small Molecules In Vivo. Chemistry & Biology 10, 313–317 (2003).

43. Hosseini-Kharat, M., Bremmell, K. E. & Prestidge, C. A. Why do lipid nanoparticles target the liver? Understanding of biodistribution and liver-specific tropism. Molecular Therapy Methods & Clinical Development 33, 101436 (2025).

44. Mui, B. L. et al. Influence of Polyethylene Glycol Lipid Desorption Rates on Pharmacokinetics and Pharmacodynamics of siRNA Lipid Nanoparticles. Molecular Therapy - Nucleic Acids 2, e139 (2013).

45. Shi, D., Toyonaga, S. & Anderson, D. G. *In Vivo* RNA Delivery to Hematopoietic Stem and Progenitor Cells *via* Targeted Lipid Nanoparticles. Nano Lett. 23, 2938–2944 (2023).

46. Fang, E. et al. Advances in COVID-19 mRNA vaccine development. Sig Transduct Target Ther 7, 94 (2022).

47. Sweeney, M. D., Sagare, A. P. & Zlokovic, B. V. Blood–brain barrier breakdown in Alzheimer disease and other neurodegenerative disorders. Nat Rev Neurol 14, 133–150 (2018).

48. Gray, M. T. & Woulfe, J. M. Striatal Blood–Brain Barrier Permeability in Parkinson’S Disease. J Cereb Blood Flow Metab 35, 747–750 (2015).

49. Zlokovic, B. V. Neurovascular pathways to neurodegeneration in Alzheimer’s disease and other disorders. Nat Rev Neurosci 12, 723–738 (2011).

50. Osaki, T. et al. miR126-mediated alteration of vascular integrity in Rett syndrome. Mol Psychiatry 10.1038/s41380-026-03492-9 (2026) doi:10.1038/s41380-026-03492-9.

51. Chen, Y., et al. Development of Highly Efficient Dual-AAV Split Adenosine Base Editor for In Vivo Gene Therapy. Small Methods 4, 2000309 (2020).

52. Iwai, H., Züger, S., Jin, J. & Tam, P.-H. Highly efficient protein *trans*-splicing by a naturally split DnaE intein from *Nostoc punctiforme*. FEBS Letters 580, 1853–1858 (2006).

53. Doench, J. G. et al. Optimized sgRNA design to maximize activity and minimize off-target effects of CRISPR-Cas9. Nat Biotechnol 34, 184–191 (2016).

54. Sanson, K. R. et al. Optimized libraries for CRISPR-Cas9 genetic screens with multiple modalities. Nat Commun 9, 5416 (2018).

55. Challis, R. C. et al. Systemic AAV vectors for widespread and targeted gene delivery in rodents. Nat Protoc 14, 379–414 (2019).

56. Wu, Y., Kirillov, A., Massa, F., Lo, W.-Y. & Girshick, R. Detectron2. (2019).

57. Stirling, D. R. et al. CellProfiler 4: improvements in speed, utility and usability. BMC Bioinformatics 22, 433 (2021).

58. Chen, D. et al. Rapid Discovery of Potent siRNA-Containing Lipid Nanoparticles Enabled by Controlled Microfluidic Formulation. J. Am. Chem. Soc. 134, 6948–6951 (2012).

