## Supplementary Information for "CHARIOT-AAV: Conjugation of diverse vectors to adeno-associated viruses for delivery of large genes"

### Supplementary Methods

#### *Cloning of split SpCas9*

To create split SpCas9, the *Rma*[37], *Npu*[35,47], and *Cfa*[36] DnaE split intein genes were synthesized (IDT, gBlocks Gene Fragments). We first removed the AAV ITR sequence from the pX330 plasmid by restriction enzyme digestion (NotI and SbfI) followed by ligation with a bridge duplex oligo, since PCR amplification with Q5 Hot Start High-Fidelity 2X Master Mix (NEB #M0494S) failed at the AAV ITR region. N-terminal split SpCas9 (from M1 to E573) including 3xFLAG and SV40 NLS and C-terminal split SpCas9 (from C574 to D1368) including nucleoplasmin NLS were amplified from the AAV ITR-removed pX330 by PCR with KOD Hot Start Master Mix (Sigma-Aldrich #71842) and assembled with split intein gene fragments by Gibson assembly (NEBuilder HiFi DNA Assembly Master Mix). Although the resulting plasmids contained correct split Cas9 and split intein sequences, the CBh promoter region harbored numerous errors. Therefore, the split Cas9 and split intein cassettes were subcloned into the original pX330 plasmid (lacking AAV ITR) by restriction enzyme digestion (EcoRI and AgeI). All gene editing assays were performed with these split SpCas9 constructs in the pX330 backbone so that both full-length and split constructs shared the same backbone. Split SpCas9 constructs were subsequently subcloned into the pAAV backbone (derived from C-Terminal Split Cas9 with GyrA intein) by restriction enzyme digestion (AgeI and EcoRI).

#### *Cloning of split dSpCas9-KRAB-MeCP2*

Split dCas9-KRAB-MeCP2 constructs were generated from dCas9-KRAB-MeCP2 (Addgene #110821). dCas9-KRAB-MeCP2 was split at several sites (L833/S834, Q844/S845, D850/S851, R859/S860, K867/S868, P872/S873, and L909/S910) by PCR.  Rma dnaE split intein sequences were amplified from the split SpCas9 (*Rma*) constructs by PCR using Q5 Hot Start High-Fidelity

2X Master Mix (NEB #M0494S). For the C-terminal construct, the CMV promoter and $dCas9_C$-KRAB-MeCP2 were separately amplified from dCas9-KRAB-MeCP2. The backbone was linearized from C-Terminal Split Cas9 with GyrA intein by restriction enzyme digestion (XbaI and SacI). The CMV promoter and split $intein_N$ were joined by overlap extension PCR. The resulting three fragments were assembled by Gibson assembly. For the N-terminal construct, the CMV promoter, NLS, and split $dSpCas9_N$ were amplified from dCas9-KRAB-MeCP2. The backbone was linearized from C-Terminal Split Cas9 with GyrA intein by restriction enzyme digestion (EcoRI and XbaI). The resulting five fragments including split $intein_N$ were assembled by Gibson assembly.

**Table S1| New constructs produced in this study**

| Name |
|---|
| pAAV-U6-sgRNA (BbsI)-CBh-NLS-Split SpCas9 (N)-Rma DnaE Intein (N)-polyA |
| pAAV-U6-sgRNA (BbsI)-CBh- Rma DnaE Intein (C)-Split SpCas9 (C)-NLS-polyA |
| pAAV-U6-sgRNA (BbsI)-CBh-NLS-Split SpCas9 (N)-Npu DnaE Intein (N)-polyA |
| pAAV-U6-sgRNA (BbsI)-CBh-Npu DnaE Intein (C)-Split SpCas9 (C)-NLS-polyA |
| pAAV-U6-sgRNA (BbsI)-CBh-NLS-Split SpCas9 (N)-Npu DnaE Intein (N)-polyA |
| pAAV-U6-sgRNA (BbsI)-CBh-Npu DnaE Intein (C)-Split SpCas9 (C)-NLS-polyA |
| pAAV-U6-sgRNA (CAG#5)-CMV-NLS-Split dSpCas9 (N; 833)-Rma DnaE Intein (N)-polyA |
| pAAV-U6-sgRNA (CAG#2)-CMV-Rma DnaE Intein (C)-Split dSpCas9 (C; 833)-NLS-KRAB-NLS-MeCP2-polyA |
| pAAV-U6-sgRNA (CAG#5)-CMV-NLS-Split dSpCas9 (N; 844)-Rma DnaE Intein (N)-polyA |
| pAAV-U6-sgRNA (CAG#2)-CMV-Rma DnaE Intein (C)-Split dSpCas9 (C; 844)-NLS-KRAB-NLS-MeCP2-polyA |
| pAAV-U6-sgRNA (CAG#5)-CMV-NLS-Split dSpCas9 (N; 850)-Rma DnaE Intein (N)-polyA |
| pAAV-U6-sgRNA (CAG#2)-CMV-Rma DnaE Intein (C)-Split dSpCas9 (C; 850)-NLS-KRAB-NLS-MeCP2-polyA |
| pAAV-U6-sgRNA (CAG#5)-CMV-NLS-Split dSpCas9 (N; 859)-Rma DnaE Intein (N)-polyA |
| pAAV-U6-sgRNA (CAG#2)-CMV-Rma DnaE Intein (C)-Split dSpCas9 (C; 859)-NLS-KRAB-NLS-MeCP2-polyA |
| pAAV-U6-sgRNA (CAG#5)-CMV-NLS-Split dSpCas9 (N; 867)-Rma DnaE Intein (N)-polyA |
| pAAV-U6-sgRNA (CAG#2)-CMV-Rma DnaE Intein (C)-Split dSpCas9 (C; 867)-NLS-KRAB-NLS-MeCP2-polyA |
| pAAV-U6-sgRNA (CAG#5)-CMV-NLS-Split dSpCas9 (N; 872)-Rma DnaE Intein (N)-polyA |
| pAAV-U6-sgRNA (CAG#2)-CMV-Rma DnaE Intein (C)-Split dSpCas9 (C; 872)-NLS-KRAB-NLS-MeCP2-polyA |
| pAAV-U6-sgRNA (CAG#5)-CMV-NLS-Split dSpCas9 (N; 909)-Rma DnaE Intein (N)-polyA |
| pAAV-U6-sgRNA (CAG#2)-CMV-Rma DnaE Intein (C)-Split dSpCas9 (C; 909)-NLS-KRAB-NLS-MeCP2-polyA |

**Table S2| Primers used in this study**

| Name | Comment |
|---|---|
| Bridge duplex oligo for pX330 to remove AAV ITR_fwd | ggccgctctccattgcgatgtgtgcgcctgca |
| bridge duplex oligo for pX330 to remove AAV ITR_rev | ggcgcacacatcgcaatggagagc |
| pX330_Cfa-C_fwd | acggcctggtggccagcaac tgcttcgactccgtggaaat |
| pX330_Cfa-C_rev | catggtggcggcctgcagaaacc ggtccaacctgaaaaaaagt |
| pX330_Cfa-N_fwd | agcaggtggacggcctgccctaa aagaattcctagagctcgctgatca |
| pX330_Cfa-N_rev | tcggtgtcgtagctcag gcactcgattttcttgaagtagtcct |
| pX330_Npu-C_fwd | aatggctttatcgccagcaat tgcttcgactccgtggaaat |
| pX330_Npu-C_rev | catggtggcggcctgcagaaaccggtccaacctgaaaaaaagt |
| pX330_Npu-N_fwd | gagtggataacctgcctaattaagaattcctagagctcgctgatca |
| pX330_Npu-N_rev | tctgtctcgtaggacaggcactcgattttcttgaagtagtcct |
| pX330_Rma-C_fwd | acgatattattgcgcataac tgcttcgactccgtggaaatc |
| pX330_Rma-C_rev | ccatggtggcggcattataataccggtccaacctgaaaaaaagt |
| pX330_Rma-N_fwd | ggagaatccctactgcctcc taagaattcctagagctcgctgatc |
| pX330_Rma-N_rev | tgagagtatcgccagccaga actcgattttcttgaagtagtcctct |
| CMV for split dCas9_C_fwd | atggctctag acgggccagatatacgcgtt |
| CMV for split dCas9_C _rev | agctctgcttatatagacctcccacc |
| Rma-C for split dCas9_fwd | tatataagcagagct ggtacc gttggaccggtattataatgccgc |
| Rma-C for split dCas9_rev | gttatgcgcaataatatcgttcgccac |
| splitdCas9_C_rev | tcagcgagctc ctatgagactctctcagtcacgggt |
| splitdCas9_C_834_fwd | attattgcgcataa ctccgactacgacgtggctg |
| splitdCas9_C_845_fwd | attattgcgcataac tcttttctcaaagatgattctattgataataaagtgttgacaa |
| splitdCas9_C_851_fwd | attattgcgcataac tctattgataataaagtgttgacaagatccgataaagctag |
| splitdCas9_C_860_fwd | attattgcgcataac tccgataaagctagagggaagagtga |
| splitdCas9_C_867_fwd | attattgcgcataac agtgataacgtcccctcagaagaagt |
| splitdCas9_C_872_fwd | attattgcgcataac tcagaagaagttgtcaagaaaatgaaaaattattggcg |
| splitdCas9_C_909_fwd | attattgcgcataac tctgagttggataaagccggct |
| CMV for split dCas9_N_fwd | aattctgcagacaaatggct atggctctag acgggccagatatacgcgtt |
| CMV for split dCas9_N _rev | atagtccatggtggc accgg tcactaaacgagctctgcttatatagacc |
| SV40 NLS for split dCas9_C_fwd | gccaccatggactataaggac |
| SV40 NLS for split dCas9_C _rev | gtacttcttgtcggctgctg |
| Rma-N for split dCas9_fwd | tgtctggctggcgatactct |
| Rma-N for split dCas9_rev | gctgatcagcgagctctagg |
| Cas9m4_845-N_fwd | gccgacaagaagtactccattgggctcgct |
| splitdCas9_N_845_rev | atcgccagccagac actggggcacgatagcagcc |
| splitdCas9_N_851_rev | atcgccagccagac aatcatctttgagaaaagactggggc |
| splitdCas9_N_860_rev | atcgccagccagac atcttgtcaacactttattatcaatagaatcatctttgag |
| splitdCas9_N_867_rev | atcgccagccagaca cttccctctagctttatcggatcttg |
| splitdCas9_N_872_rev | atcgccagccagac aggggacgttatcactcttcc |
| splitdCas9_N_909_rev | atcgccagccagac acaggccacctcgttcagcc |

**Table S3| sgRNA sequences used in this study**

| Name | Comment |
|---|---|
| sgRNA_StopLight_fwd | CACCG GGACAGTACTCCGCTCGAGT |
| sgRNA_StopLight_rev | AAAC ACTCGAGCGGAGTACTGTCC C |
| sgRNA_CAG_2_fwd | CACCG AAGCGCTAATTACAGCCCGG |
| sgRNA_CAG_2_rev | AAAC CCGGGCTGTAATTAGCGCTT C |
| sgRNA_CAG_5_fwd | CACCG ACTGACCGCGTTACTCCCAC |
| sgRNA_CAG_5_rev | AAAC GTGGGAGTAACGCGGTCAGT C |
| sgRNA_CAG_6_fwd | CACCG TCGGAGCGCGCCGCTCTGAT |
| sgRNA_CAG_6_rev | AAAC ATCAGAGCGGCGCGCTCCGA C |
| sgRNA_CAG_7_fwd | CACCG GCTCACCTGTGGGAGTAACG |
| sgRNA_CAG_7_rev | AAAC CGTTACTCCCACAGGTGAGC C |
| sgRNA_Ai14_fwd | CACCG AAGTAAAACCTCTACAAATG |
| sgRNA_Ai14_rev | AAAC CATTTGTAGAGGTTTTACTT C |

### Supplementary Data

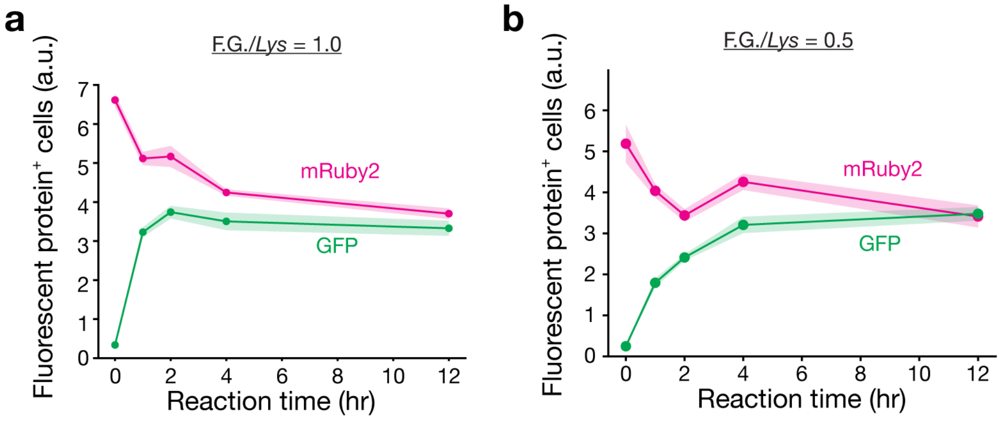


**Fig. S1| Experimental study of conjugation chemistry. a**, **b**, Percentage of HEK293T cells that expressed GFP or mRuby2. HEK293T cells were transduced with AAV-AAV, which were prepared with varying conjugation times. AAV-DJ-CAG-mRuby2 and AAV9-CAG-NLS-GFP were employed. The number ratio of functional groups (F.G.; i.e. Tetrazine (Tz) and *trans*-cyclooctene (TCO)) to exposed *Lys* residues on AAV capsids was 1.0 (**a**) or 0.5 (**b**). At a F.G.-to-*Lys* ratio of 1.0, GFP expression peaked at the 2-hour reaction time. In contrast, GFP expression did not saturate even after 12 hours at a F.G./*Lys* ratio of 0.5. mRuby2 expression decreased as conjugation reaction time. Data are represented as mean ± standard error of the mean (S.E.M.), n = 3 biological replicates.

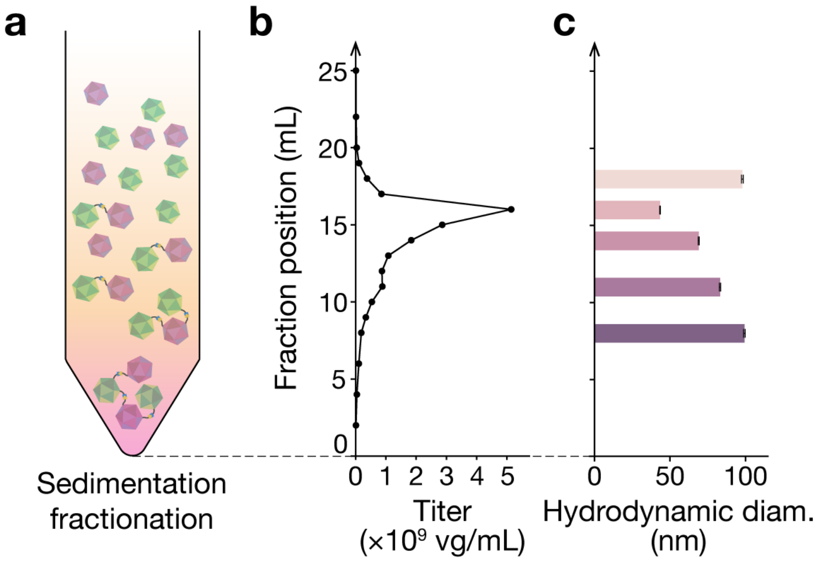


**Fig. S2| Fractionation of AAV-AAV conjugates by ultracentrifugal sedimentation. a**, Schematic of gradient ultracentrifugation. A sucrose gradient was prepared (see Methods), and ~200 µL of AAV-AAV conjugate solution was placed on top of the lightest layer. The tube was spun at 150,000 ×g for 1.5 hours at 4 °C. After centrifugation, every 1 mL of the solution was collected from the bottom of the tube using a syringe and a needle. **b**, Titer distribution of the fractionated AAV-AAV conjugate solution. The fractionated 1 mL solutions were directly subjected to quantitative polymerase chain reaction (qPCR) measurement to determine the titers. **c**, Hydrodynamic diameter of AAV-AAV conjugates purified from different fractions. The fractionated AAV-AAV conjugates in sucrose solution were medium-exchanged with Dulbecco's phosphate buffered saline with 0.1% Pluronic F-68 (DPBS-F68) using filter centrifuge units to achieve the same viscosity across samples. Bars and error bars represent mean ± standard deviations (S.D.), n = 8 independent measurements.

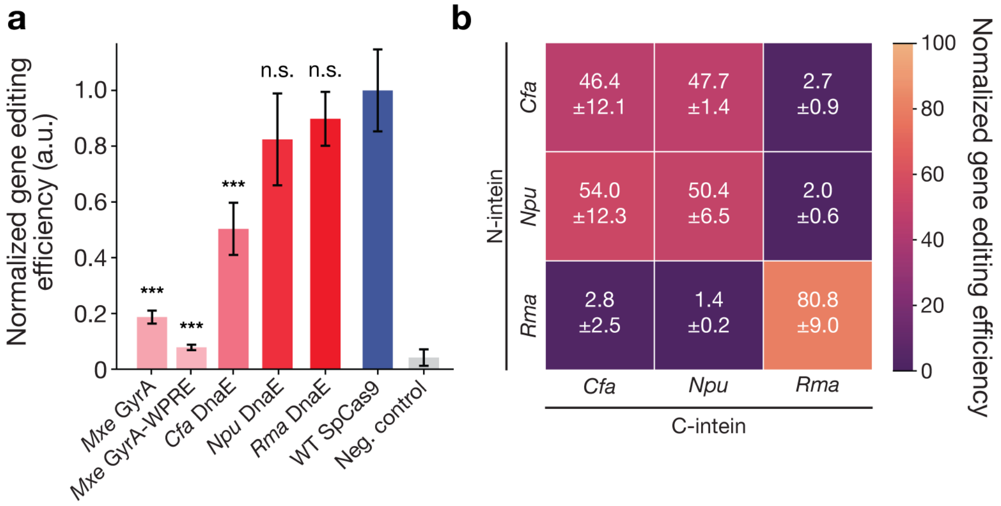


**Fig. S3| Evaluation of split inteins from different organisms and host proteins. a**, Split SpCas9 constructs with diverse split inteins were evaluated by lipofection and compared with full-length SpCas9. StopLight HEK cells were employed in this assay. Inteins from *Rma* DnaE and *Npu* DnaE showed comparable gene editing activity to the full-length control. **b**, Cross-reactivity between DnaE inteins from different organisms. Inteins from *Npu* and *Cfa* exhibited substantial cross-reactivity, while *Rma* DnaE inteins did not, suggesting that the *Npu* and *Rma* DnaE intein pair could serve as orthogonal inteins for a three-split design. Bars and error bars represent mean ± S.E.M. Statistical significance was assessed by Welch's t-test (n = 3 biological replicates).

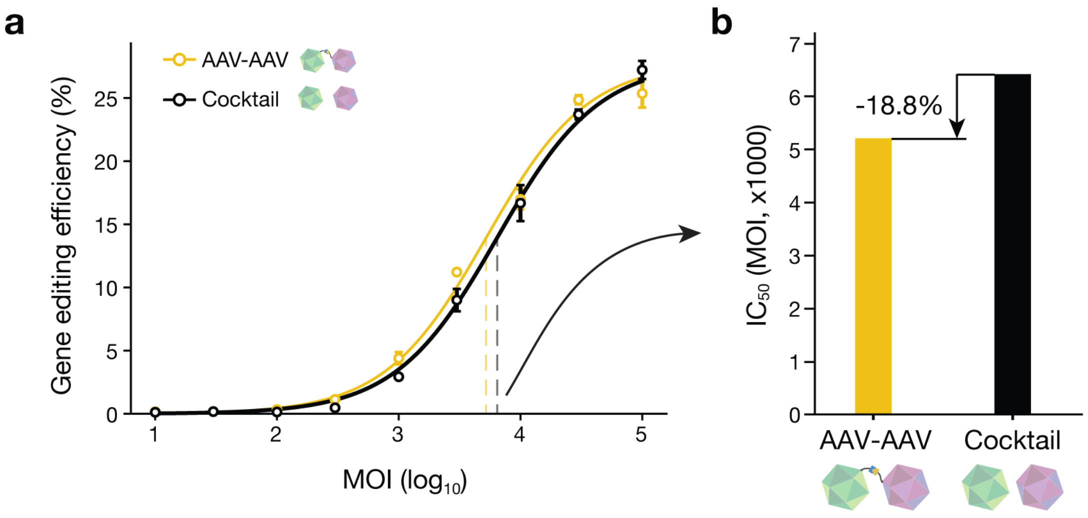


**Fig. S4| IC50 analysis of AAV-AAV conjugates in StopLight HEK cells.** **a**, Gene editing efficiency in StopLight HEK cells transduced with AAV-AAV conjugates or cocktails, both packaging the split SpCas9 constructs. **b**, $IC_{50}$ values derived from the dose-response curves in **a**. AAV-AAV conjugates exhibited an 18.8% lower $IC_{50}$, indicating that an 18.8% lower dose is sufficient to achieve the same gene editing efficacy. Data are presented as mean ± S.E.M. (n = 3).

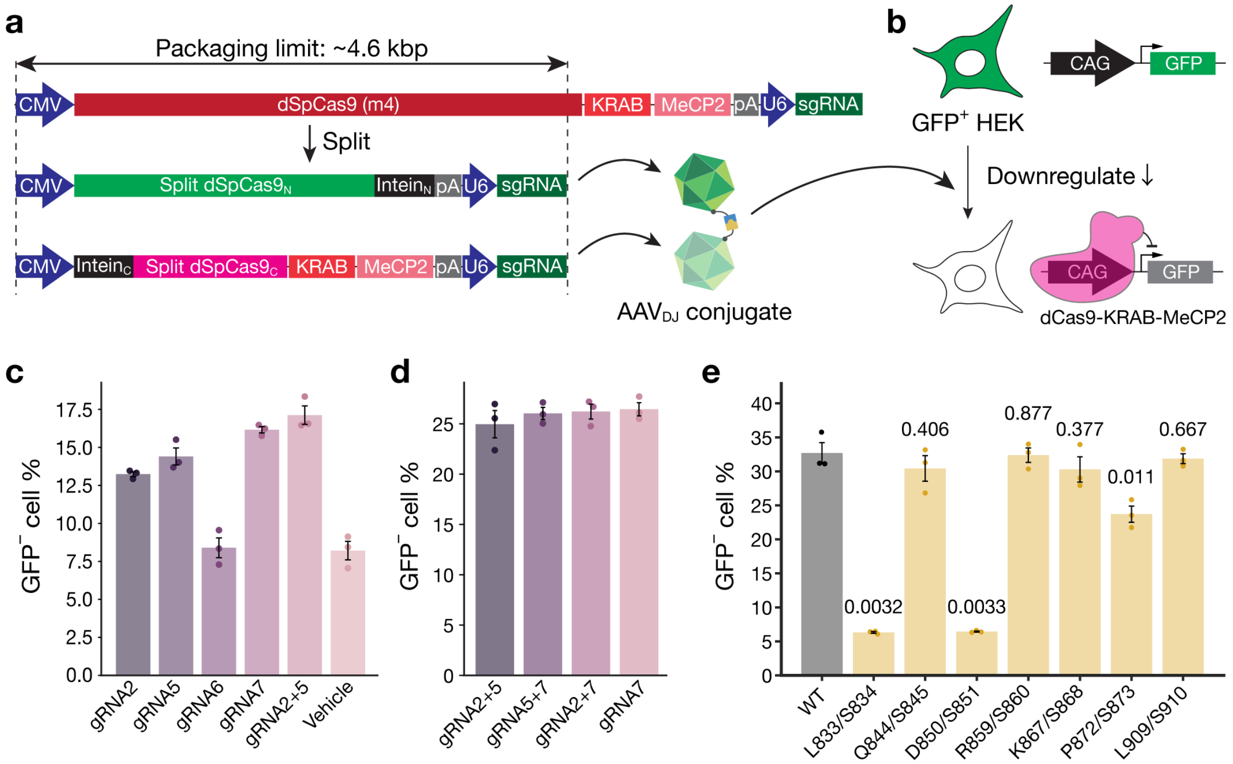


**Fig. S5| Design of split dSpCas9-KRAB-MeCP2. a**, Schematic of split dSpCas9-KRAB-MeCP2 construct design. dSpCas9 (dCas9m4) was split at different sites and tested for activity. Since the full construct is longer than the SpCas9 gene editing cassette, the split site needed to be shifted toward the C-terminus to accommodate the sgRNA cassette in both split constructs. **b**, Schematic of in vitro gene repression experiments. GFP-expressing HEK cells were employed in this assay, which expressed GFP driven by the CAG promoter. sgRNAs targeted the transcription start site of the CAG promoter, resulting in GFP downregulation after successful delivery. **c**, **d**, Different sgRNA target sites were tested in GFP-expressing HEK cells. At 72 hours after lipofection with full-length dSpCas9-KRAB-MeCP2 and sgRNAs, GFP levels were analyzed by flow cytometry. Since two different sgRNA sequences can be loaded into the two split constructs, mixtures of two sgRNAs were also evaluated. sgRNA #2 and #5 were selected for the main constructs. **e**, Different split sites within dSpCas9 (dCas9me) were tested, and their gene repression efficiency was compared to that of full-length dSpCas9-KRAB-MeCP2. Multiple split sites exhibited equivalent gene repression activity relative to the full-length control. As L909/910 was closest to the middle of the genetic construct among constructs that exhibited comparable activities to the full-length control, this split site was selected. Bars and error bars represent mean ± S.E.M. Statistical significance was assessed by Welch's t-test (n = 3 biological replicates).

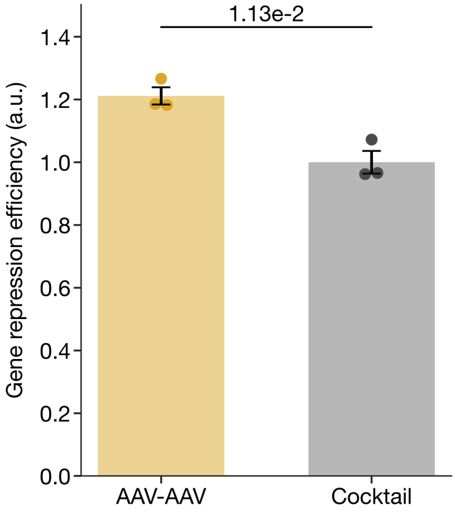


**Fig. S6| In vitro evaluation of AAV-AAV conjugate-mediated epigenetic gene repression by split dSpCas9-KRAB-MeCP2.** The selected split dSpCas9-KRAB-MeCP2 constructs were packaged in AAV-DJ capsids, which were then conjugated and fractionated. GFP-expressing HEK cells were transduced with the resulting AAV-AAV conjugates or cocktails. At 48 hours post-transduction, cells were analyzed for GFP levels by flow cytometry. AAV-AAV conjugates exhibited 1.21-fold greater gene-repression relative to the cocktail. Bars and error bars represent mean ± S.E.M. Statistical significance was assessed by Welch's t-test (n = 3 biological replicates).

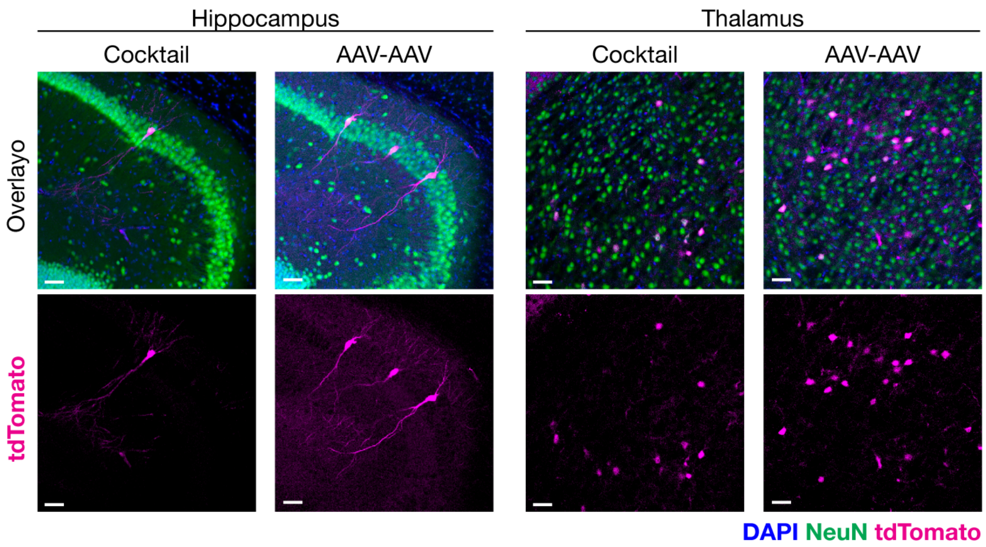


**Fig. S7| In vivo evaluation of AAV-AAV conjugate-mediated gene editing with split SpCas9 in Ai14 mice.** Representative confocal images of the hippocampus and thalamus from mice intravenously injected with AAV$_{\text{CAP-B10}}$-AAV$_{\text{CAP-B10}}$ conjugates or AAV cocktails ($2.5\times10^{10}$ vg/animal). Scale bars, 50 µm.

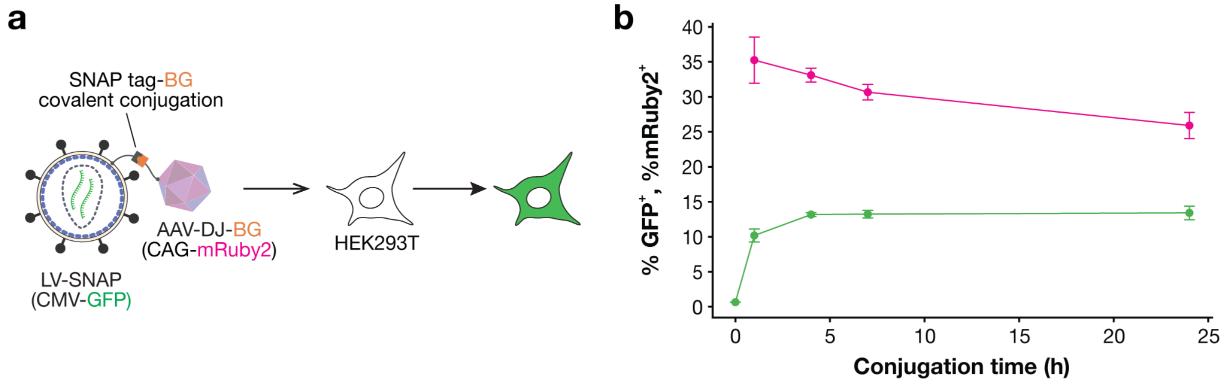


**Fig. S8| Reaction time optimization for AAV-LVV conjugation. a**, Schematic of the in vitro evaluation. Plasmids encoding mutated VSV-G and SNAP-tag were employed for LVV production. pLenti-CMV-GFP was packaged in the LVV. AAV-DJ-CAG-mRuby2 was functionalized with $O^6$-benzylguanine (BG) via NHS chemistry. The functionalized particles were conjugated for varying durations. At defined time points, BG-mPEG24 was added to quench the SNAP-tag and incubated at 4 °C for 4 hours. The resulting AAV-LVV conjugates were directly used for transduction of HEK293T cells. At 24 hours post-transduction, cells were analyzed for GFP expression by flow cytometry. **b**, Percentage of GFP-positive and mRuby2-positive cells transduced with AAV-LVV conjugates produced at different conjugation times. The GFP-positive cell fraction plateaued at conjugation times longer than 4 hours, while the mRuby2-positive cell fraction decreased continuously with conjugation time, indicating over-conjugation of AAV-DJ. Data are presented as mean ± S.E.M (n = 3),

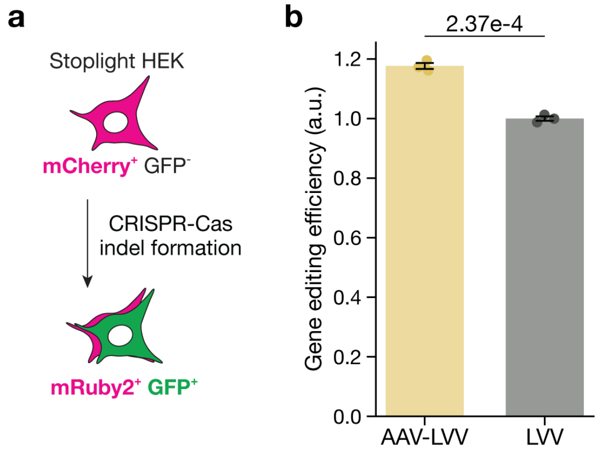


**Fig. S9| Gene editing by SpCas9 delivered via AAV-LVV in vitro. a**, StopLight HEK cells were used in this assay. Upon gene editing at the target site, GFP expression was induced. **b**, Gene editing efficiency of AAV-LVV conjugates and cocktails in StopLight HEK cells at 24 hours post-transduction. AAV-DJ was conjugated with LVV particles. LVVs were loaded with SpCas9-based gene editing cassettes with control sgRNA scaffold (lentiCRISPR v2; Addgene #52961), while AAV-DJ packaged sgRNA for StopLight HEK. Bars and error bars represent mean ± S.E.M. Statistical significance was assessed by Welch's t-test (n = 3 biological replicates).

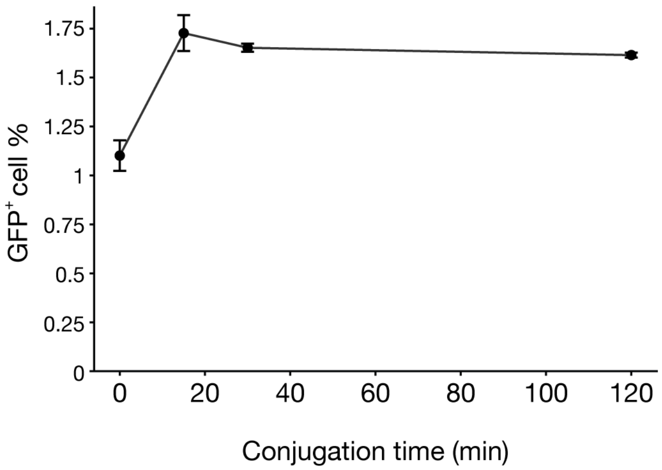


**Fig. S10| Reaction time optimization for AAV-LNP conjugation.** AAV-TCO and LNP-Tz were conjugated via the IEDDA click chemistry, followed by quenching. The resulting AAV-LNP conjugates were evaluated for transfection efficiency of the LNP component (GFP mRNA) in HEK293T cells. To prevent non-specific transfection, HEK293T cells were incubated in serum-free medium during transfection (see Methods; LNP/cell ratio = 3,160). At 24 hours post-transfection, cells were analyzed for GFP expression by flow cytometry. Data are represented as mean ± S.E.M. (n = 3).

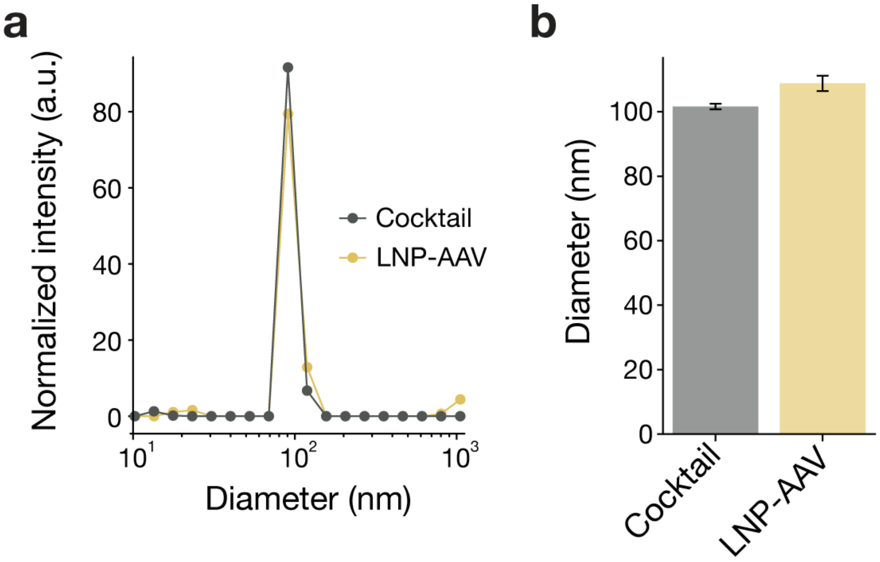


**Fig. S11| Dynamic light scattering analysis of AAV-LNP conjugates. a**, Hydrodynnamic diameter distribution of AAV-LNP conjugates and the cocktail measured by dynamic light scattering (DLS) in DPBS-F68 at 23 °C. A polydisperse distribution was assumed. **b**, Mean hydrodynamic diameters of the cocktail and AAV-LNP conjugates. Bars and error bars represent mean ± S.D., n = 8 independent measurements.

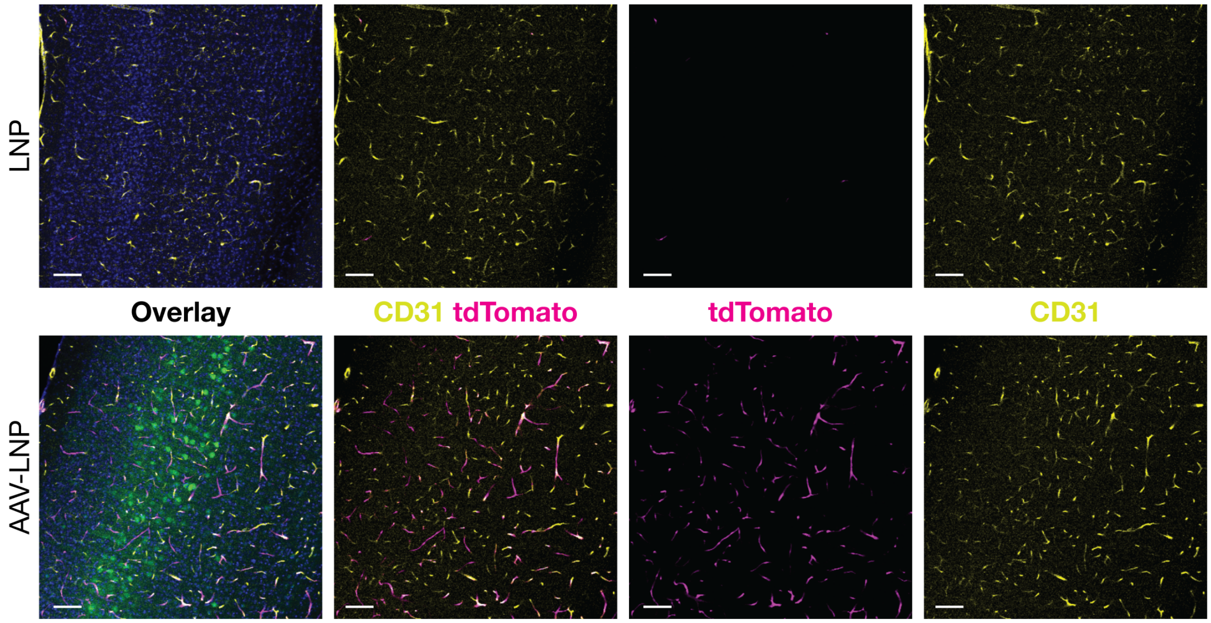


**Fig. S12| Cortical brain vasculature from AAV$_{CAP\text{-}B10}$-LNP-injected mice.** Sagittal brain sections from mice injected with either AAV$_{CAP\text{-}B10}$-LNP conjugates or LNP controls were stained with a vasculature marker CD31. LNPs packaged Cre recombinase mRNA and AAV$_{CAP\text{-}B10}$ packaged pAAV-CAG::mNeonGreen. Five days post-injection, the Ai14 mice were sacrificed and perfused with 4% PFA. CD31 was stained by immunohistochemistry (Methods). Quantification of tdTomato fluorescence intensity in CD31$^{+}$ objects was performed using CellProfiler4 (Fig. 3i). Blue – DAPI, Green – mNeonGreen, Magenta – tdTomato, Yellow – CD31. Scale bars, 100 µm.

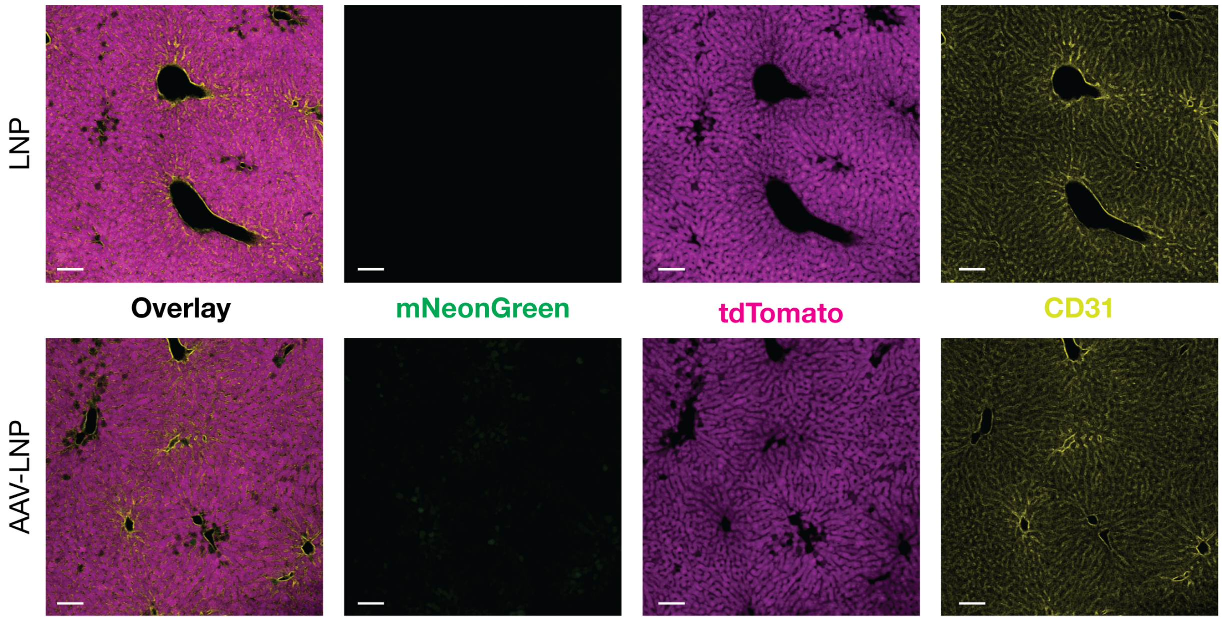


**Fig. S13| Confocal images of the liver from AAV$_{\text{CAP-B10}}$-LNP-injected mice.** Liver tissues were sectioned at 50 µm thickness and stained with CD31 via immunohistochemistry (Methods). Ai14 mice were systemically administered AAV$_{\text{CAP-B10}}$-LNP or LNP. AAV$_{\text{CAP-B10}}$ packaged pAAV-CAG::mNeonGreen and LNPs packaged Cre recombinase mRNA. Five days post-injection, the animals were sacrificed and perfused with 4% PFA. Green – mNeonGreen, Magenta – tdTomato, Yellow – CD31. Scale bars, 100 µm.

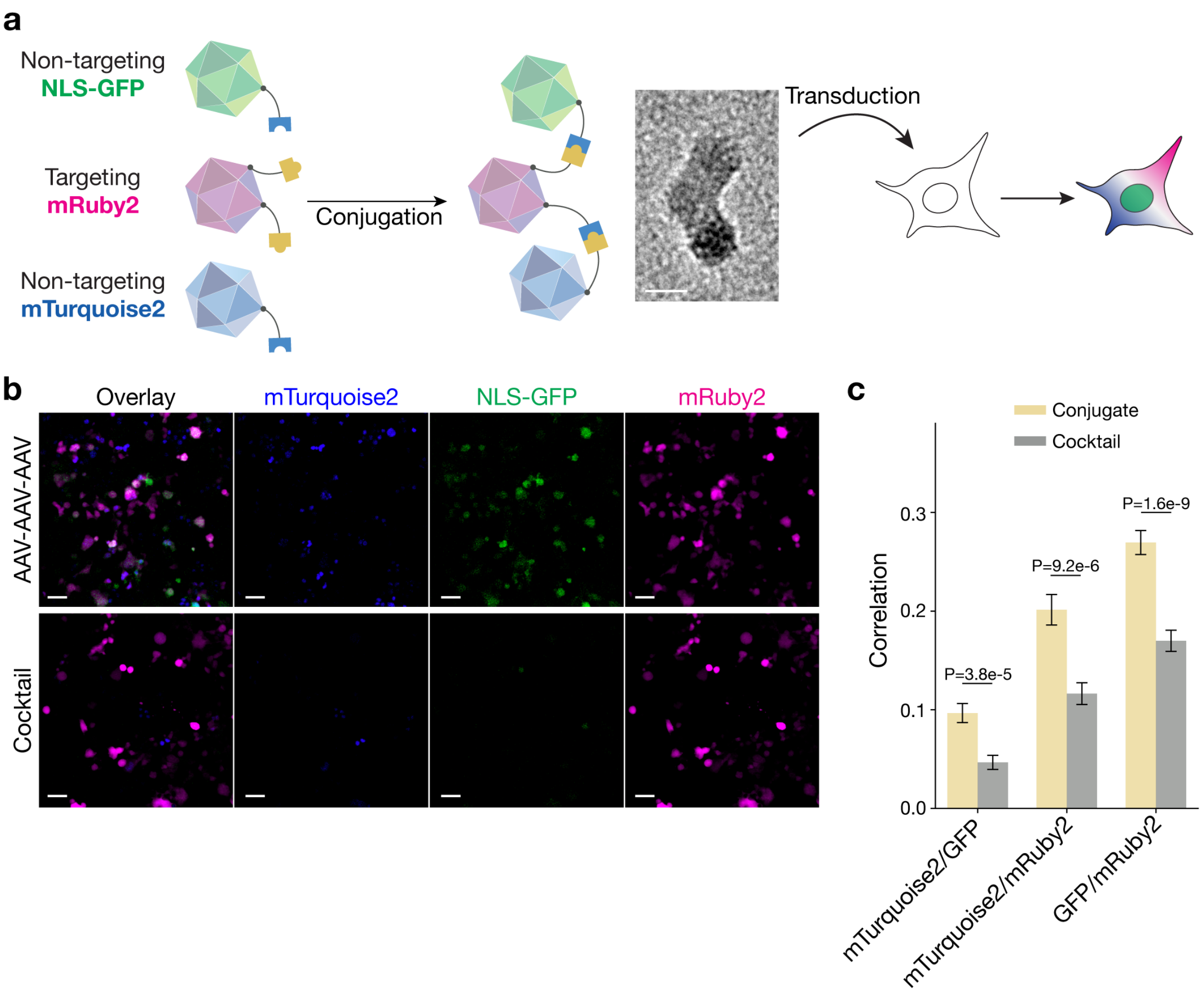


**Fig. S14| Proof-of-concept experiments with three-AAV conjugates. a**, Schematic of the three-AAV conjugation experiment. AAV-DJ and AAV9 served as the targeting and non-targeting serotypes, respectively. AAV-DJ packaged CAG-mRuby2, while two different batches of AAV9 packaged either CAG-NLS-GFP or CAG-mTurquoise2. AAV-DJ was functionalized with Tz, and AAV9 capsids with TCO. The functionalized capsids were then mixed and conjugated via click chemistry. The produced AAV-AAV-AAV conjugates were tested in HEK293T cells. A cocktail of intact AAVs (from the same batches) served as a control. Scale bar, 20 nm. **b**, Confocal images of HEK293T cells transduced with AAV-AAV-AAV conjugates or the cocktail. At 24 hours post-transduction, cells were imaged on a confocal microscope (Leica Stellaris 5 ) at 37 °C under 5% $CO_2$. Scale bars, 50 µm. **c**, Pearson's correlation coefficients among fluorescence signals of the three fluorescent proteins. Quantification was performed using CellProfiler. Bar plot represents mean ± S.E.M (n = 353 for AAV conjugates, n = 304 for AAV cocktails).
